# An Integrated Decarbonising Approach to Mitigate Methane Emissions and Enhance Productivity in Rice Cultivation

**DOI:** 10.1101/2025.10.17.683017

**Authors:** Kins Varghese, Ali Ma, Thankaraj Salammal Maria Shibu, Kasthurirengan Sampath, Kenny J.X. Lau, Zhongchao Yin, Naweed I. Naqvi, Srinivasan Ramachandran

## Abstract

Flooded rice paddies contribute approximately 12% of global anthropogenic methane emissions, accounting for 1.5% of the total warming effect from all greenhouse gases. With the rising global demand for rice due to population growth, the need for effective methane mitigation strategies in rice cultivation is increasingly critical. This study investigates the combined impact of irrigation methods, fertiliser combinations, and varietal differences on productivity, water use and methane emissions in rice. Field trials were conducted across five land parcels covering 8 Ha in the Sathyamangalam region of Tamil Nadu, India, from October 2024 to January 2025. Results revealed that drip irrigation significantly reduced seasonal methane emissions by up to 68% (128 kg/ha/season) compared to continuous flood irrigation (402.32 kg/ha/season) offering a sustainable solution to address climate change. Furthermore, our modified package of practices coupled with tailored fertiliser combination, led to a 28% reduction in methane emissions (222 kg/ha/season) relative to continuous flooding methods used by the farmers in the region (309 kg/ha/season). Methane emission differences of up to 23.8% were also evident across rice varieties ADT-45 (229.7 kg/ha/season and BPT-5204 (301.7 kg/ha/season) for. Although flood irrigation yielded a 5–6% higher grain productivity than drip irrigation, the TLL fertiliser package under flood irrigation still provided distinct benefits with a yield increase of 5.4%. Notably, water usage under drip irrigation was 42.5% lower on average across the five locations, with minimal impact on yield, resulting in a marked improvement in water use efficiency (0.62 kg.m^-3^ under drip vs. 0.39 kg.m^-3^ under flood irrigation). Our findings highlight the value of integrating modern irrigation techniques, optimizing fertiliser management, and appropriate varietal selection with higher environmental sustainability and improved farm productivity to mitigate climate impact in rice cultivation.

## Introduction

Rice (Oryza sativa), the staple food for more than half the global population, is a significant source of anthropogenic greenhouse gas (GHG) emissions, contributing 6–11% of global methane (CH_4_) emissions. Methane, with a global warming potential 28 times that of CO_2_, is primarily generated through anaerobic decomposition by methanogenic archaea in continuously flooded paddy soils (Conrad, 2007). Annual methane emissions from rice fields are estimated at 24–31 teragrams, accompanied by 130 gigagrams of nitrous oxide N_2_O) (IPCC, 2021; FAO, 2022; Cui et al., 2021). With global rice demand projected to increase by 114 million tonnes by 2035 (Suzanne et al., 2012), these exacerbate emissions accelerate climate change, which in turn poses threats to crop yield and food security. Methane emissions in rice cultivation are influenced by multiple factors, including soil properties, climatic conditions, and crop management practices such as irrigation, varietal selection, and fertiliser application (Yagi et al., 1997; Wassmann et al., 2000; Bharati et al., 2001).

The traditional continuous flood (CF) provides agronomic benefits such as weed suppression and yield stability, but it creates anaerobic soil conditions leading to elevated methane emissions (Neue, 1993), contributing to global warming and indirectly affecting crop productivity and soil health. Continuous flooding (CF reduces redox potential (∼ −150 mV), which fostering an anaerobic environment conducive to methanogenesis with approximately

90% of methane emitted to the atmosphere via the aerenchyma tissues of rice plants (Conrad, 1996). Mid-season drainage, introduces oxygen into the soil, thereby suppressing methanogenesis and promotes methanotroph activity (Conrad, 1996). Methane emission from flooded fields vary widely, typically ranging from 20 to 100 kg CH_4_ per hectare per season, depending on soil type, organic inputs, and climatic conditions (Yan et al., 2009). Alternative irrigation strategies including intermittent irrigation, alternate wetting and drying (AWD), and aerobic rice systems (Hafsa et al., 2020; Price et al., 2013) and drip irrigation (Parthasarthi et al., 2019) has shown to reduce methane emissions by enhancing soil aeration and disrupting anaerobic conditions. Drip irrigation, in particular support sustainable rice production while mitigating greenhouse gas emissions without compromising yield (Wassmann et al., 1993; Theivasigamani et al., 2019). Studies reported that drip irrigation can reduces methane emissions by 70–80%, N_₂_O by ∼34%, and total global warming potential (GWP) by 25–44% while using 23.3% less water, increasing yields by 15–23%, and improving water productivity by 2–4 times compared to conventional flooding (Parthasarathi et al., 2019; Mallareddy et al., 2023,; Demirel et al., 2020). These findings highlight the potential of drip irrigation method as a water-efficient and environmentally sustainable method for rice cultivation, particularly in water-scarce regions (Ramulu et al., 2016; Sharda et al., 2017; Padmaja & Malla Reddy, 2018; Natarajan et al., 2020). Despite occasional yield reductions, substantial water savings make drip irrigation a viable alternative to conventional flooding.

Methane emissions from rice fields vary substantially among cultivars due to differences in root architecture, biomass, and aerenchyma development. Varieties with extensive aerenchyma facilitate methane transport from soil to atmosphere, whereas those with limited aerenchyma and reduced root exudation restrict emissions (Qin et al., 2015). High-yielding varieties are often associated with lower emissions (Malyan et al., 2016; Chen et al., 2021) and some cultivars with higher methane emissions tend to support more active methanogenic communities, while those with reduced emissions may limit microbial activity in the rhizosphere (Hu et al., 2023). These patterns can vary across growth stages and are not always consistent (Kim et al., 2018). Modern breeding of shorter-duration rice varieties may contribute to reduction in cumulative methane emissions. Furthermore, recently developed cultivars enhance methane oxidation through increased root exudation and radial oxygen loss, underscoring the critical role of cultivar selection in sustainable, low-emission rice production (Li et al., 2022).

The combination of NPK fertilisers with secondary nutrients and micronutrients has significantly enhance rice productivity while simultaneously mitigating methane emissions. Gypsum is recognized for its role in improving soil structure and the nutrient availability, thereby which increasing the yields. Long-term application of gypsum has been reported to enhance nutrient dynamics on rice farms in Southern India (Prakash et al., 2025) Meng et al. (2023) demonstrated that gypsum application increased rice yield by 58%, reduced methane emissions by 47%, GWP by 22%, and greenhouse gas intensity (GHGI) by 31%, without elevating N_2_O emissions. The incorporation of ferrous sulphate (FeSO_4_) has been associated with improved soil conditions, contributing to a reduction in methane emissions (Khatun et al., 2020). Zinc enhances root development, increase soil redox potential, and supports methane-oxidizing bacteria, resulting in a 39.57% yield increase under basal application (Shikha et al., 2022) and increases productive tillers compared to foliar application (Qaisrani & Saeed, 2011). Boron application has been reported to increase rice yield by 25% when applied at the flowering stage and up to 34% when applied at both tillering and flowering stages (Bodeerath et al., 2024). Magnesium sulphate (MgSO4) improves photosynthetic efficiency and root zone aeration, indirectly contributing to methane mitigation.

Supplementation with sulphate- and iron-containing fertilisers, such as CaSO4 and FeSO4, alters the soil environment to favour sulphate- and iron-reducing bacteria. These microbial groups compete with methanogens by utilizing shared substrates, including hydrogen _and_ acetate, thereby suppressing methanogenesis through competitive exclusion (Kristjansson et al., 1982; Denier et al., 2001). Iron-rich amendments have been shown to effectively reduce methane emissions from organic matter inputs, such as rice straw, by stimulating methanotrophic activity (Hu et al., 2020; Schütz et al., 1989; Jakobsen et al., 1981; Achtnich et al., 1995). Alternative electron acceptors, including nitrate (NO_3-_), ferric iron (Fe^3+^), manganese (Mn^4+^), and sulphate (SO_4_^2-^) have been reported to suppress methane formation by inhibiting methanogens. Although sulfate additions do not eliminate methane production, they can reduce emissions by up to 35% with degree of suppression dependent on acetate and sulphate concentrations (Minamikawa et al., 2015, Sinke et al., 1992; Gupta et al., 1994). Despite these insights, few studies have examined the combined effect of fertiliser and water management on greenhouse gas emission in rice (Della et al., 2024; Islam et al., 2022), and there is a lack of comprehensive research on integrated influence rice cultivars, irrigation, and fertiliser on methane mitigation. To address this critical knowledge gap, the present study evaluated the interactions of two local rice varieties (ADT 45 and BPT 5204) with irrigation methods [continuous flooding (CF) and drip irrigation (DI)], fertiliser regimes (optimized TLL recommended dosage fertiliser and traditional farmer’s practices) and water use efficiencies, the objective of advancing decarbonize rice production through reduced methane emission.

## Materials and Methods

### Experiment site and climatic conditions

Field trial sites located in and around the Sathyamangalam region of Tamil Nadu, India. Geographical locations and co-ordinates of trial sites are given in the Table 1. Field trials were conducted in five sites encompassing a total area of 8 Ha (Figure 1). Soil textures across the sites ranged between clay loam to sandy clay loam. Soil pH varied between 7.7 and 8.3, indicating neutral to slightly alkaline conditions. The soil organic carbon content ranged between 0.8% to 1.2%, it reflects moderate level of soil organic matter across the trial locations (Table 2).

**Figure 1:**
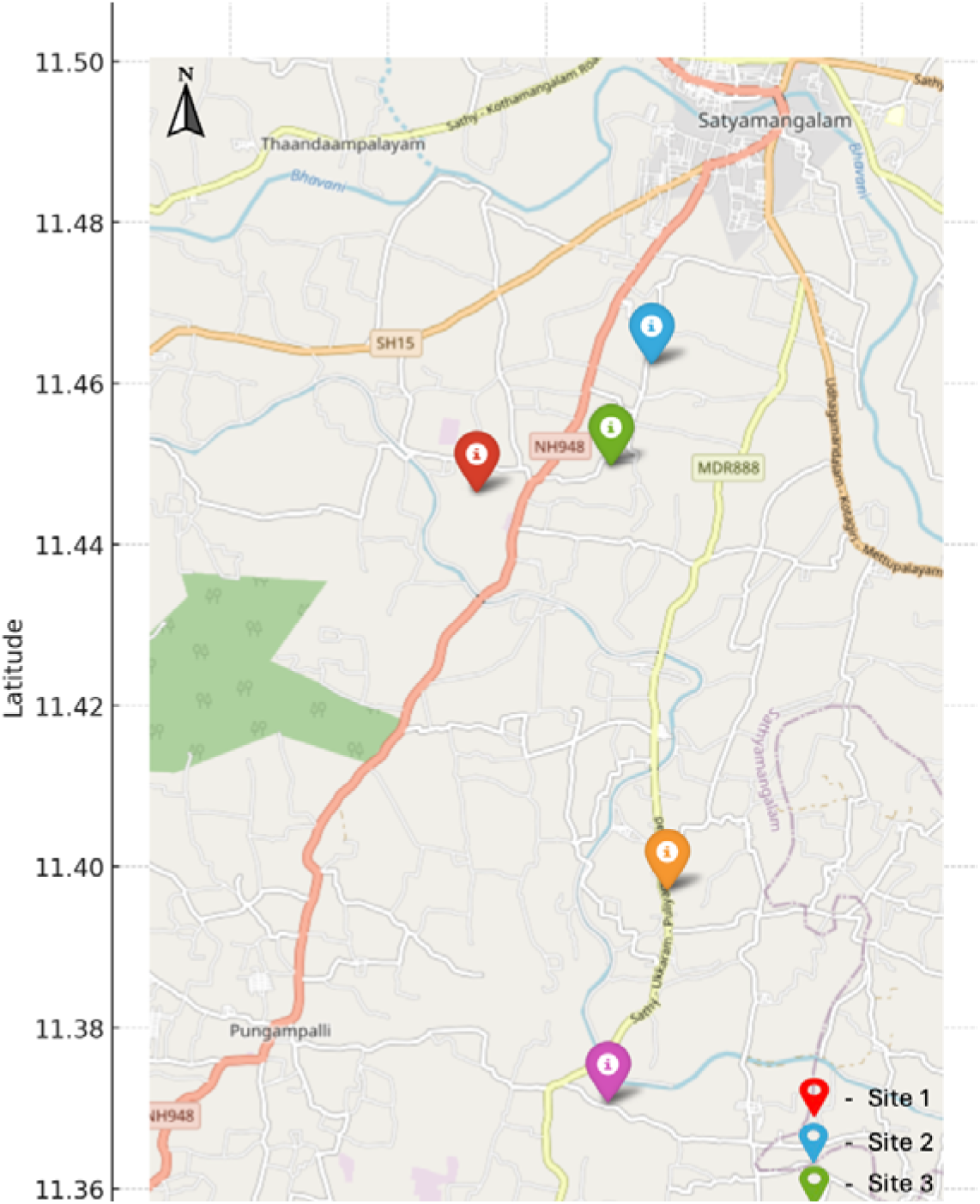
Experimental sites in Sathyamangalam region of Tamil Nadu, India.

**Table 1:** Trial sites, area and co-ordinates in Sathyamangalam, Tamil Nadu, India.

| Sites | Area (in Ha) | Co-ordinates |
| --- | --- | --- |
| Site 1 | 0.61 | 11°27'38.91" N, 77°12'45.62" E |
| Site 2 | 1.21 | 11°27'50.49" N, 77°13'37.31" E |
| Site 3 | 1.62 | 11°23'44.13" N, 77°13'35.30" E |
| Site 4 | 1.82 | 11°25'8.46" N, 77°14'2.02" E |
| Site 5 | 2.83 | 11°28'28.21" N, 77°13'53.75" E |

**Table 2:**
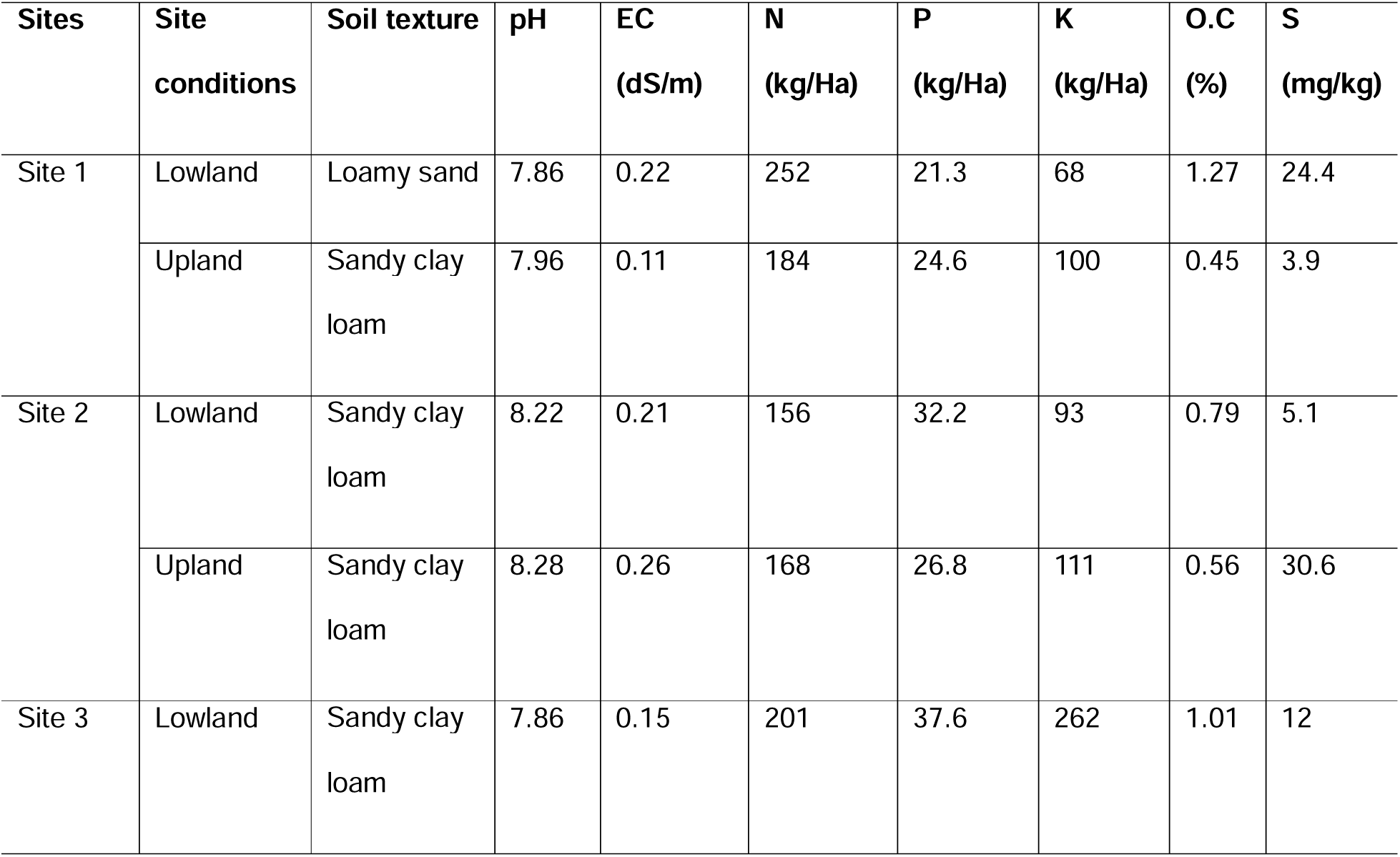
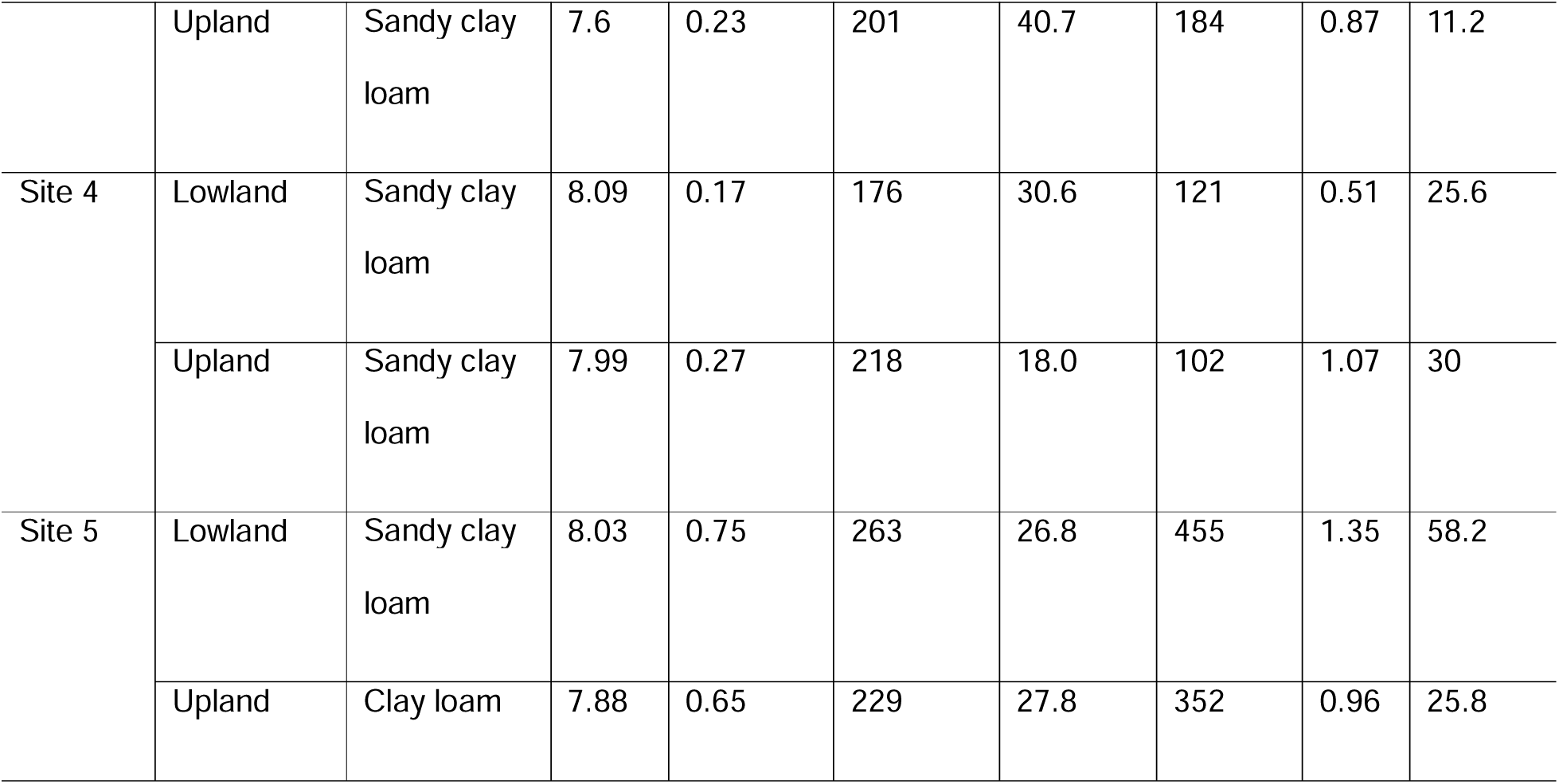
Physio-chemical properties of soil at five trial sites in Sathyamangalam, Tamil Nadu, India before initiation of different treatment trials.

The trial site designed to study the effects of rice varieties (ADT 45 and BPT 5204), two irrigation methods (continuous flood and drip irrigation), fertiliser combinations (TLL-F and FP) on the productivity and methane emissions reduction, and also to evaluate water usage efficiency (WUE). Current trial region has a tropical climate with distinct wet and dry seasons that strongly influence agricultural practices. In this region, rice is the predominant staple crop, traditionally cultivated under canal irrigation with conventional fertiliser compositions. Trial sites receive an average annual rainfall of approximately 630 mm with mean temperatures ranging from 27°C to 30°C. Rice is cultivated during two principal growing seasons that overlap with regional climatic patterns and water availability.

During the trial period (October 2024–January 2025), field experiments were conducted on five sites covering 8 Ha and involving eight small holders from Sathyamangalam region of Tamil Nadu, India. The fields were managed by local farmers under the technical supervision of Temasek Life Sciences Laboratory (TLL), to ensure uniformity and adherence to operating procedures. Across five trial sites, an average of 196.5 mm of rainfall was recorded, and average temperatures ranged between 26°C and 28°C. These management practices and environmental conditions were focused to evaluating the treatment effects on rice performance.

### Trial details and experiment design

The trial was designed to evaluate the effects of three factors each at 2 levels: irrigation method (Continuous Flood (CF) and Drip Irrigation (DI), fertiliser combinations-TLL fertiliser mix (TLL-F), and Farmer practice (FP) and rice varieties-ADT-45 (V1) and BPT-5204 (V2). This experimental setup was structured using Split-split plot design with irrigations in the main plot, fertilisers application at subplot and varieties allocated to the sub-sub plots. This design allowed us to evaluate the interactions between the three factors at multiple levels, while the block design minimized variability within the experimental plots by grouping them based on similar characteristics.

The trial includes 8 treatment combinations in four replications, each site with 32 plots (totally 160 plots across 5 sites) (T1-CF+V1+TLL-F, T2-CF+V2+TLL-F, T3-CF+V1+FP, T4-CF+V2+FP, T5-DI+V1+TLL-F, T6-DI+V2+TLL-F, T7-DI+V1+FP, T8-DI+V2+FP). In the main plot, two irrigation methods were adopted to supply water to the plants. The first method was drip-irrigation, where scheduling was based on the temperature and reference evapotranspiration (ETo) rates. ETo was estimated using pan evapotranspiration data and Manna software (Rivulis Irrigation Systems, Israel). The second method was continuous flood irrigation, where a standing water level of 3–5 cm was maintained throughout the growing period. Water usage was recorded using a water meter installed at the irrigation outlet. Intermittent rainfall contributed additionally to the water supply during the cropping period which was monitored using rain gauges at each site.

In sub-plot, two fertiliser combinations were tested, i) TLL-F, a modified fertiliser combinations developed and tested by Temasek Lifesciences Laboratory (TLL), Singapore specifically for rice GHG emission reduction, ii) region specific general farmer practice (FP). Fertilisers under TLL-F applications were done according to the package of practices developed by TLL and whereas FP were applied as per the farmers’ schedule, A detailed fertiliser dosages and combination, and application regime were outlined in table 3. For sub-sub plot, two rice varieties were placed for the trials. ADT-45 a medium duration, semi-dwarf variety developed by Tamil Nadu Rice Research Institute (TNRRI), Aduthurai and BPT-5204, a semi dwarf, medium to long duration variety developed by Indian Agricultural Research Institute (IARI), India. Before transplanting on to the plots, seedlings were raised in a nursery bed for 21-days after sowing.

**Table 3:** Fertilization application rate (kg/ha) and schedule for five trial sites.

|  | Basal fertilizer |  |  |  |  | Tillering |  |  |  | Panicle Initiation |  | Heading |
| --- | --- | --- | --- | --- | --- | --- | --- | --- | --- | --- | --- | --- |
|  | N | P<sub>2</sub>O<sub>5</sub> | K<sub>2</sub>O | CaSO<sub>4</sub> | FeSO<sub>4</sub> | N | ZnSO<sub>4</sub> | MgSO<sub>4</sub> | Boron | N | K<sub>2</sub>O | N |
| TLL-RDF | 40 | 50 | 25 | 250 | 30 | 30 | 20 | 20 | 2 | 30 | 25 | 30 |
| FP | 60 | 75 | 60 |  |  | 60 |  |  |  | 60 |  |  |

### Collection and measurements of methane emissions

Methane emissions from the trial plots were collected systematically using the closed chamber method, a widely accepted technique for field-scale greenhouse gas measurement (Minamikawa et al., 2015). Two chambers with dimensions of 50 x 50 x 65 cm made up of transparent acrylic sheets were used to collect the air samples. The base chambers were placed in each plot 24 hours prior to sample collection. Bottom edges of the chambers were sealed with soil to prevent any leakage. On the day of sample collection, the upper chamber installed with DC fan to homogenise the air were placed on top of the base chamber without disturbing the base.

Air samples were collected using syringes (20-25 ml) from the chambers through a window sealed with rubber septum. These air samples are then transferred to sealed 20 ml glass vials filled with water. The air samples injected into the sealed gas vials to displace the 18 ml of water and leaving approximately 2 ml of water. Air sampling was carried out at weekly intervals throughout the rice growing season to capture temporal variations in methane emission patterns. Sampling was conducted during the daytime, between 8:30 a.m. and 11:30 a.m., a period known to provide fluxes representative of daily averages. At each sampling event, gas samples were collected from the chambers at four time points: immediately upon chamber closure (0 minutes), and subsequently at 10, 20, and 30 minutes. This sequential sampling allowed the calculation of methane fluxes by assessing concentration changes over time. Analysis was performed using a Gas Chromatography with Flame Ionization Detector (GC-FID) (Shimadzu Corporation), with Pora PLOT Q-HT column (CP7559, Agilent Technologies) was used to separate methane from the air samples.

### Methane hourly gas flux

Linear regression is used for calculating the hourly methane flux. This method is based on the principle that the concentration gradient of methane between flooded soil and the atmosphere is quite large, so that methane is emitted at a constant rate. The hourly fluxes of methane (mg CH_4_ m^−2^ h^−1^) is calculated as follows:

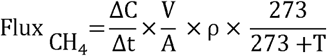

where ΔC/Δt is the concentration change over time (ppm-CH_4_ h^−1^); V - chamber volume (m^3^); A - chamber area (footprint; m^2^); ρ - gas density (0.717 kg m^−3^ for methane at 0°C); and T - the mean air temperature inside the chamber (°C).

### Estimation of Rice Productivity, water usage, and Greenhouse Gas Emissions (GHG)

To evaluate the actual impact of different irrigation methodologies, fertiliser combinations, and rice varieties on crop yield, production measurements were conducted till maturation stage. The analysis involved measuring the economic output from replications, ensuring that the data were representative of typical field conditions. These productivity measurements allowed for a detailed comparison of the various treatments and their effects on yield.

### Yield estimation

Harvested grains were dried to a moisture content of 12-13%. Measured 1,000 grains weight and yield per plot was used to estimate the yield per hectare, considered 7-8% yield loss while extrapolating the yield per plot data to yield per Ha.

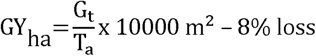

where,

G_t_= grain yield per plot (at grain moisture content of 12-13%%), T_a_ = plot area (m^2^), GY_ha_= estimated grain yield per hectare.

### Global warning potential (GWP) of methane

The Global Warming Potential (GWP) of methane (CH_₄_) emissions was calculated as carbon dioxide equivalents (CO_2_ ^e^) over a 100-year time horizon, using the radiative forcing potential of 28 for CH_4_ relative to CO_₂_, as outlined in the IPCC article 6 guidelines (2013). This approach enabled the comparison of GHG emissions across different treatments, facilitating an understanding of the environmental impact of various irrigation and fertilization practices.

### Statistical data analysis

Statistical analyses were carried out using Excel and SPSS version 29. Each treatment was conducted at split-spilt design with replicates. The mean differences were carried out using ANOVA analysis and Tukey HSD test, and Least Significant Difference (LSD) was calculated at the P<0.05 and P<0.01 level. Experiments were carried out in 4 replications to ensure repeatability and to understand the effects of treatments carried out on crop’s productivity and methane emission reduction.

## Results and Discussion

Growth performance of rice varieties across sites:

Growth and developmental patterns of two rice varieties were largely similar across the five experimental sites. Significant differences were observed in plant height and the number of effective tillers per plant across irrigation treatments, indicating the influence of irrigation on plant growth. The maximum plant height was recorded in treatment T4 (95.3 cm), followed by treatment T6 (94.7 cm) across all sites (Figure. 2). Averaged across eight treatments and five sites, continuous flooding produced taller plants (93.5 cm), compared to DI (91.6 cm) (Supplemental Table 1). Similarly, the number of effective tillers per plant (ETPP) and grains per panicle (GPP) were significantly affected by both irrigation method and varietal selection. Treatment T4 resulted in highest ETPP (8.1 plant^-1^) followed by T2 (7.8 plant^-1^) with lowest observed in T7 (6.36 plant^-1^) (Figure 2). Across irrigation methods, continuous flooding resulted in the highest ETPP (7.6 plant^-1^), whereas DI produced the lowest (6.9 plant^-1^). Significant varietal effect (p = 0.01) was also observed, with BPT-5204 producing more ETPP (7.7 plant^-1^) than ADT-45 (6.8 plant^-1^) (Supplemental Table 1). GPP was highest with T1 (169.3), followed by T3 (167.9) and lowest with T6 (141.9). Continuous flood produced more GPP (158.5) compared to DI (149.2). Among varieties, ADT-45 produced the more GPP (162.4) than BPT-5204 (145.3) (Supplemental Table 1). Fertiliser combinations did not have a significant on ETPP and GPP across five sites.

**Figure 2:**
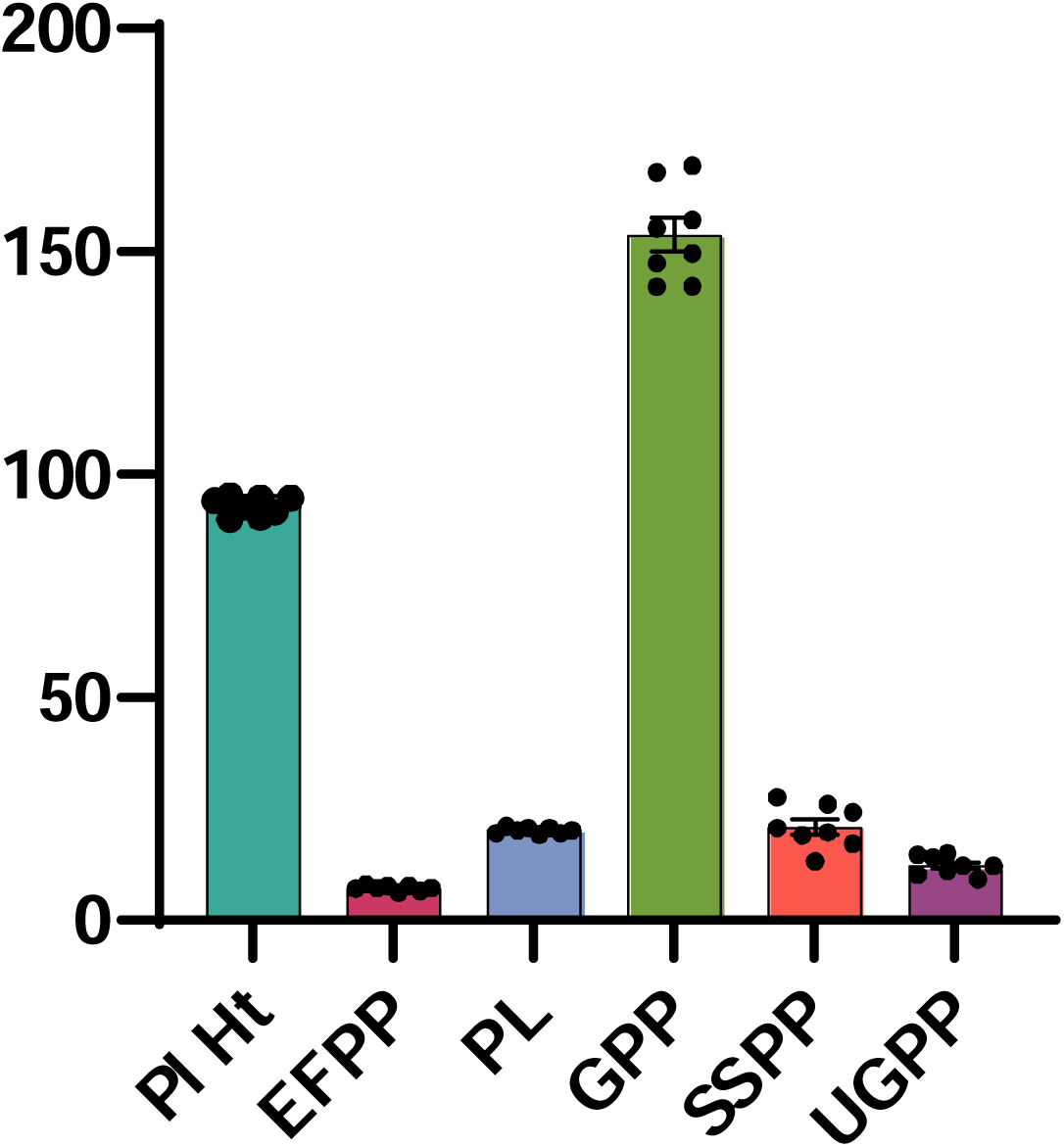
Comparative impact of eight treatments on rice plant growth performance across five sites in Sathyamangalam, Tamil Nadu, India. In the graph, each bar represents average value of a growth parameter under different treatments, “•” indicating the specific mean value for each treatment from five sites.

The observed reduction in plant height and effective tillers under drip irrigation may be attributed to increased evapotranspiration (ET) rates and limited water availability, which likely prompt plants to allocate resources toward survival rather than vegetative growth. This water stress induces morphological adjustments resulting in shorter plants and fewer tillers as a conservative adaptive response. In contrast, continuous flood irrigation provides stable soil moisture, facilitating enhanced tiller initiation and overall plant development. The sensitivity of tiller formation to water regimes has been documented in previous studies (Bhandari et al. 2003, He et al. 2022, Joshi et al. 2018, Sasmita et al. 2022, Wang et al. 2023). Moreover, Zhao et. al., (2023) highlighted the importance of varietal traits, reporting that timely irrigation combined with balanced nutrients can stabilize the grain number per panicle by optimizing resource allocation and mitigating stress effects.

Panicle length (PL) was consistently greater in the variety BPT-5204 (20.7 cm) compared to ADT-45 (19.6 cm) when averaged across sites, highlighting the influence of genotype on this trait. Among treatments, T8 (20.9 cm) exhibited the highest PL followed by T4 (20.8 cm), with lower observed in T5 (19.3 cm) (Figure 2). Differences due to irrigation methods Significant variation in panicle length was observed across fertiliser treatments (p>0.05), indicating that nutrient management exerted a measurable influence under the prevailing trial conditions, with farmer practice producing slightly higher PL (20.3 cm) compared to TLL-F (20.0 cm) (Supplemental Table 1). These results suggest that panicle length in rice is primarily determined by genetic factors, and nutrient availability, remaining relatively stable under non-stressful environmental conditions, rather than being substantially influenced by moderate variations in water availability. Previous studies have similarly shown that cultivar-specific traits are the major determinants of panicle architecture and length (Hemalatha et al. 2018, Latif et al., (2005), Thakur et al. (2011), Hossaen et al. (2011). While severe water stress during at reproductive stages can affect panicle development in some genotypes, most evidence indicates that, in non-stress environments, genetic and nutrient factors are the dominant determinants (Panda et al. 2020).

The number of sterile spikelets per panicle (SSPP) and unfilled grains/panicle (UGPP) were significantly influenced by treatments across all sites. Treatment T1 exhibited the lowest SSPP and UGPP% (13.2% and 9%, respectively) followed by T2 (73.2% and 10.2%), while the highest value was recorded in T8 (27.6% and 15 %) (Figure 3). Continuous flood significantly reduced the SSPP and UGPP% (18.6% and 10.6 %, respectively) compared to drip irrigation 23.4% and 14%. Fertiliser regimes also affected these traits with TLL-F producing lower SSPP and UGPP (17.6% and 11. 4%) compared to farmer practices (24.4% and 13.3%) respectively (Supplemental Table 1). Varietal effects were generally non-significant, except at site 3, where ADT-45 exhibited lower SSPP and UGPP (17% and 7.74%) than BPT-5204 (20.4% and 9.9%) (Supplemental Table 2). These results highlight the predominant influence of environmental factors and agronomic management on spikelet sterility in rice. Prolonged soil moisture under continuous flooding enhances spikelet fertility and reduces sterility, particularly during fertilization and grain filling, demonstrating its relative superiority over DI (Tsujimoto et al., 2021; Krishnasree et al. 2024). Additionally, the inclusion of boron and other essential micronutrients in the TLL-F blend contributed to reduction in sterile spikelet. This observation was consistent with previous studies reporting the positive effects of boron on reproductive success and zinc sulphate in spikelet fertility and overall grain yield (Pachauri et al. 2024, Maqsood et al. 1999, Khan et al., 2002, Qadir et al. 2013, Ankur et al. 2020, Prakash et al. 2015, Bu et al. 2025)

**Figure 3:**
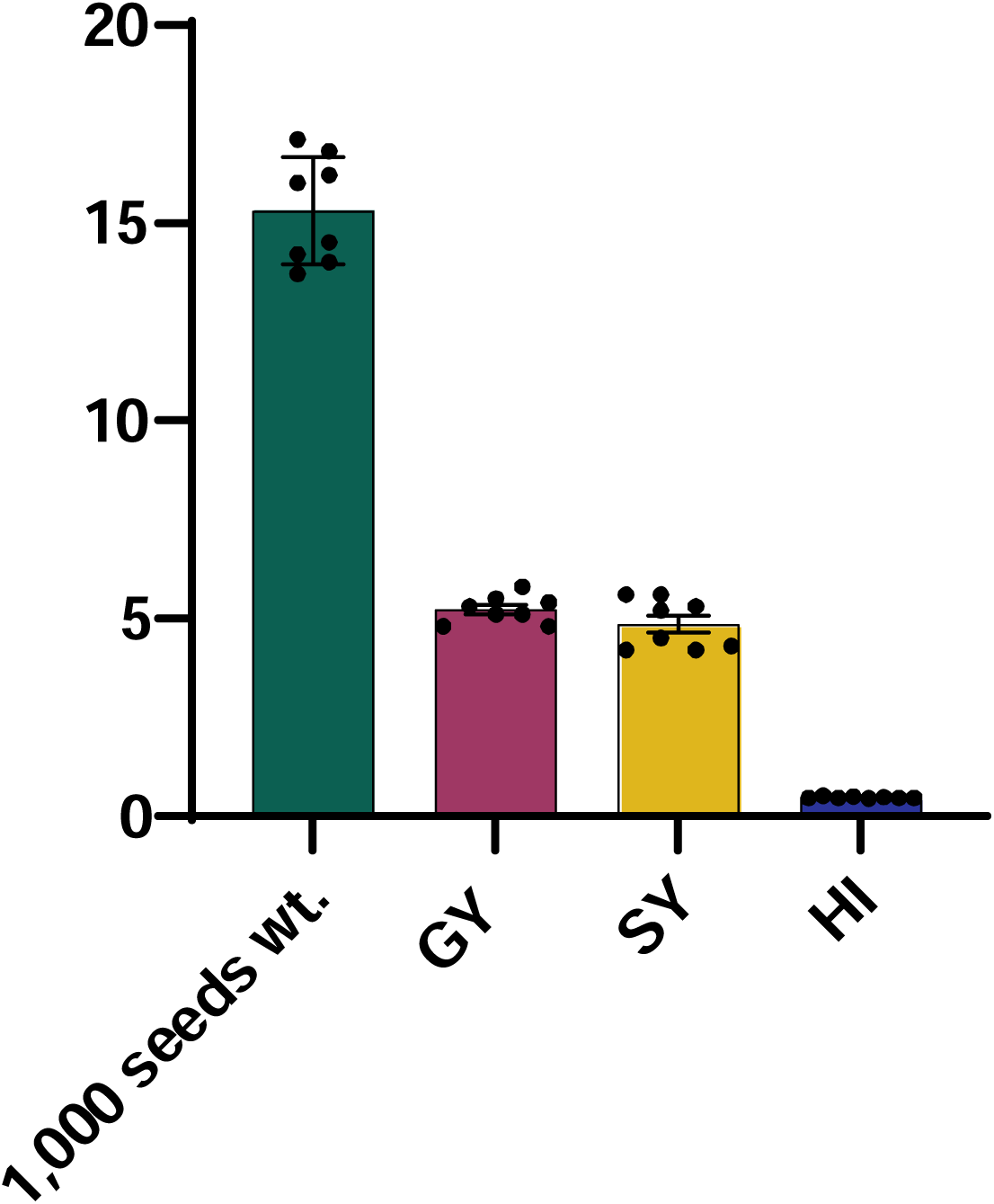
Comparative impact of eight treatments on rice yield performance across five sites in Sathyamangalam, Tamil Nadu, India. In the graph, each bar represents average value of a growth parameter under different treatments, “•” indicating the specific mean value for each treatment from five sites.

The 1,000-seed weight (TW), grain yield (GY) and harvest index (HI) were significantly influenced by irrigation regime, fertilisers management and varietal differences. Averaged across five sites, treatment T1 exhibited the highest 1,000 seed weight (17.0 g), followed by T3 (16.8 g) with lowest observed in T6 (13.8) (Figure 3). Across five sites, continuous flooding consistently resulted in higher 1,000-seed weight (15.6 g) compared to drip irrigation (15.0 g). (Supplemental Table1). Among fertiliser treatments, our fertiliser formulation (TLL-F) enriched with micronutrients produced higher 1,000-seed weight (15.4 g) relative to farmer practice (15.2 g) in site 1 and 2, with no significant differences in the remaining sites (Supplemental Table 2). Between the two varieties, ADT-45 exhibited the highest 1,000-seed weight (16.5 g), outperforming BPT-5204 (14.1 g) (Supplemental Table 1). Significant interactions were observed between variety and irrigation (V x I) as well as among variety, irrigation and fertiliser (V x I x F) for the seed test weight.

The highest grain yield, straw yield, and harvest index were recorded in treatment T2 (5.8 t/ha, 4.9 t/ha and 0.52 respectively), followed by T4 (5.5 t/ha, 4.6 t/ha and 0.5 respectively), with the lowest value observed in T5 (4.8 t/ha and 3.8 t/ha respectively) and T7 (0.46 for harvest index)(Figure 3). Continuous flood produced the maximum grain yield and harvest index (5.5 t/ha and 0.5 respectively) compared to drip irrigation. Similarly, TLL-F resulted in higher yield and harvest index (5.4 t/ha and 0.5 respectively) relative to conventional farmer practice. Among the varieties, BPT-5204 exhibited the highest grain yield, straw yield and harvest index (5.4 t/ha, 5.5 t/ha and 0.5 respectively) 5 across five sites to ADT-45 (Supplemental Table 2).

Beser et al. (2016), reported that the highest test weight in rice was achieved under continuous flood across different irrigation methods. Several field and physiological studies have demonstrated that rice grain filling and yield are strongly influenced by varietal traits and water management strategies. Continuous flood has been found to be particularly effective in promoting grain development and increasing seed weight (Cebi et. al., 2023). Other studies have highlighted the sensitivity of grain weight to water availability during grain filling, particularly when using sprinkler or drip irrigation methods as alternatives to continuous flooding (Sasmita et al., 2022). Previous studies also documented higher yields, improved tiller survival and increased aboveground biomass under continuous flooding compared to drip irrigation (Demirel et al. 2020, Vijayakumar et al. 2019, Sharma et al. 2019). In our observations, grain yield in rice is largely genotyped dependent, governed by traits such as panicle number, spikelet fertility, and grain weight. Yield variability has been well-documented, with substantial attributed to genotypic characteristics (Patak et al., 2011; Joshi et al., 2013 Laza et al. 2004).

Varietal plays a critical role in determining role in grain and straw productivity (Hari Om et al. 1997). Our observation indicates that harvest index also varies among genotypes and overall productivity is influenced by management practices, reflecting intrinsic physiological mechanisms that govern assimilate partitioning and stress tolerance (Golada & Debbarma 2023, Amanullah & Inamullah, 2016, Uzzaman et al. 2015). Bu et al. (2025) demonstrated that under continuous flooding, rice plants allocate more resources to productive growth rather than stress adaptation. However, Cebi et al. (2023) noted that yield reduction under drip irrigation can be environment-dependent, with certain conditions narrowing the performance gaps between irrigation methods. Consequently, optimizing rice production requires the strategic alignment of rice genotypes to management practices and local environmental conditions to fully exploit genotypic potential under both conventional and water-saving regimes.

The superior performance of TLL-F treatment over farmer’s practices can likely be attributed to the inclusion of micronutrients such as zinc and boron which play critical roles in enzymatic activation, photosynthetic efficiency, respiration hormonal regulation. Debnath and Ghosh (2011), and Choudhary et al. (2015), reported that zinc application enhanced grain yield by 24.52% compared to the control plots. Similar trends have been observed in other studies with yield increases of 24.8% and 37.7%, respectively (Mahata et al. 2013, Naik and Das, 2010), Notably, Debnath et al. (2015) demonstrated that the combined application of boron and zinc synergistically enhanced the yield potential.

The absence of significant interaction effects among irrigation methods, fertiliser regimes, and varietal differences indicates that these agronomic factors largely operated independently, without exhibiting synergistic or antagonistic interactions. This suggests that the effects irrigation, fertilisation, and varietal selection on growth and yield parameters were distinct and additive, rather than interdependent. Continuous flooding consistently enhanced grain filling and yield across both varieties and fertiliser treatments, regardless of their combinations. Similarly, the TLL-F treatment improved spikelet fertility and reduced the percentage of unfilled grains, irrespective varietal selection or irrigation systems.

### Water usage

Water usage (m^3^/ha) for rice cultivation under continuous flood and drip irrigation at five sites in Sathyamangalam, Tamil Nadu from October 2024 to February 2025 is summarized in Figure 4. Total rainfall during the experimental period was approximately 200 mm. Water use under drip irrigation ranged from 7,668 to 9,508 m^3^/ha, whereas continuous flood ranged from 12,523 to 16,503 m3/ha. On average, drip irrigation reduced water consumption by ∼40%, consistent with previous studies on precision irrigation in rice (Bouman et al., 2007). Maximum water savings were observed at site-2 (45.5%), likely due to improved emitter uniformity and favourable soil texture, which enhanced water retention. Site-5 exhibited the highest water use under both irrigation methods, reflecting elevated percolation losses and increased evapotranspiration demand (Supplemental Table 3). Previous studies have demonstrated that drip irrigation significantly reduced overall water consumption, highlighting its potential for sustainable water conservation and its relevance to global food security, particularly in drought prone regions. These finding corroborate previous the results of Anusha and Nagaraju. (2015), who reported a 52% reduction in water use with drip irrigation compared to continuous flooding and are supported by other studies (Parthasarathi et al., 2018, Sidhu et al., 2019 and Sasmita et al., 2022). Overall, drip irrigation represents a critical tool for the sustainable intensification of rice production while conserving valuable freshwaters resources for future generations.

**Figure 4:**
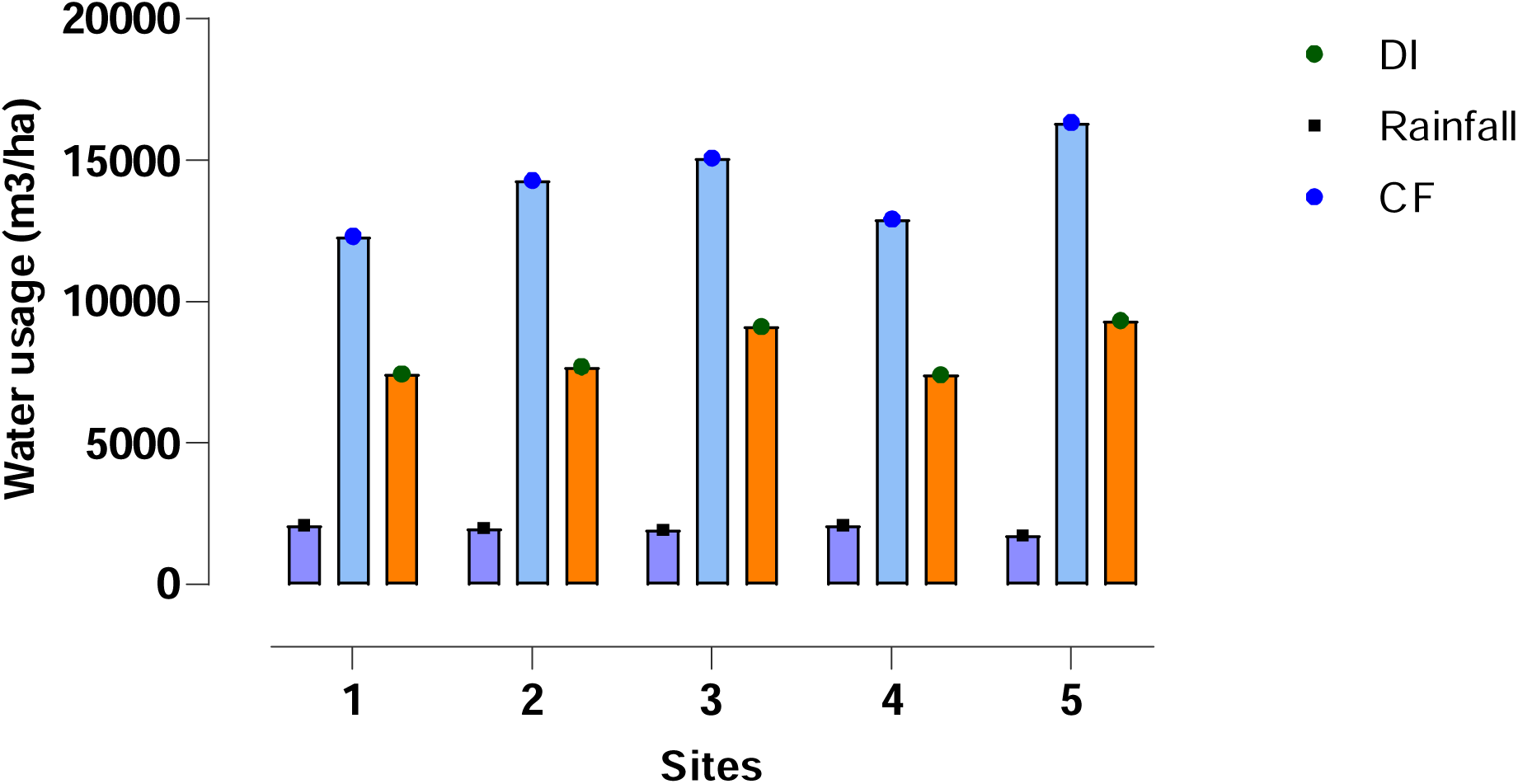
Water Usage (m^3^ ha^-1^) under continuous flooding (CF) and drip irrigation (DI) for five sites in Sathyamangalam, Tamil Nadu, India

### Water Productivity (WP)

Across all five sites, drip irrigation achieved 50–64% higher water productivity (WP) than continuous flood, mainly by reducing evaporation, percolation, and runoff, while maintaining soil moisture near field capacity. This improved root growth, nutrient uptake, and grain filling, supported by better stomatal conductance, photosynthesis, and reduced stress. Site 4 recorded the highest water productivity under drip irrigation (0.74 kg/m³) attributed to favourable soil properties and compatibility between cultivar–fertiliser, whereas Site 5 exhibited the lowest water productivity under continuous flooding (0.28 kg/m³), with drip irrigation partially improving it to 0.45 kg/m³ (Figure 5). These results consistent with previous multi-location drip irrigation trials, which reported water productivity improvements ranging from 0.46 – 0.67 kg/m^3^ across diverse rice varieties and growing conditions (Padmanabhan et al., 2019). Additional studies further affirms that the superiority of drip irrigation for enhancing water productivity (Komballi et al., 2016; He et al., 2013; Sharda et al., 2017; Soman et al.,2018). Cultivar traits and balanced fertiliser regimes, particularly those enriched with secondary and micronutrients (Ca, Mg, Zn, B), further enhance water productivity by improving nutrient-use efficiency and reducing spikelet sterility (Shahane et al., 2019; Lenka et al., 2001). Such nutrient supplementation supports improved plant growth and grain filling while minimizing water requirements.

**Figure 5:**
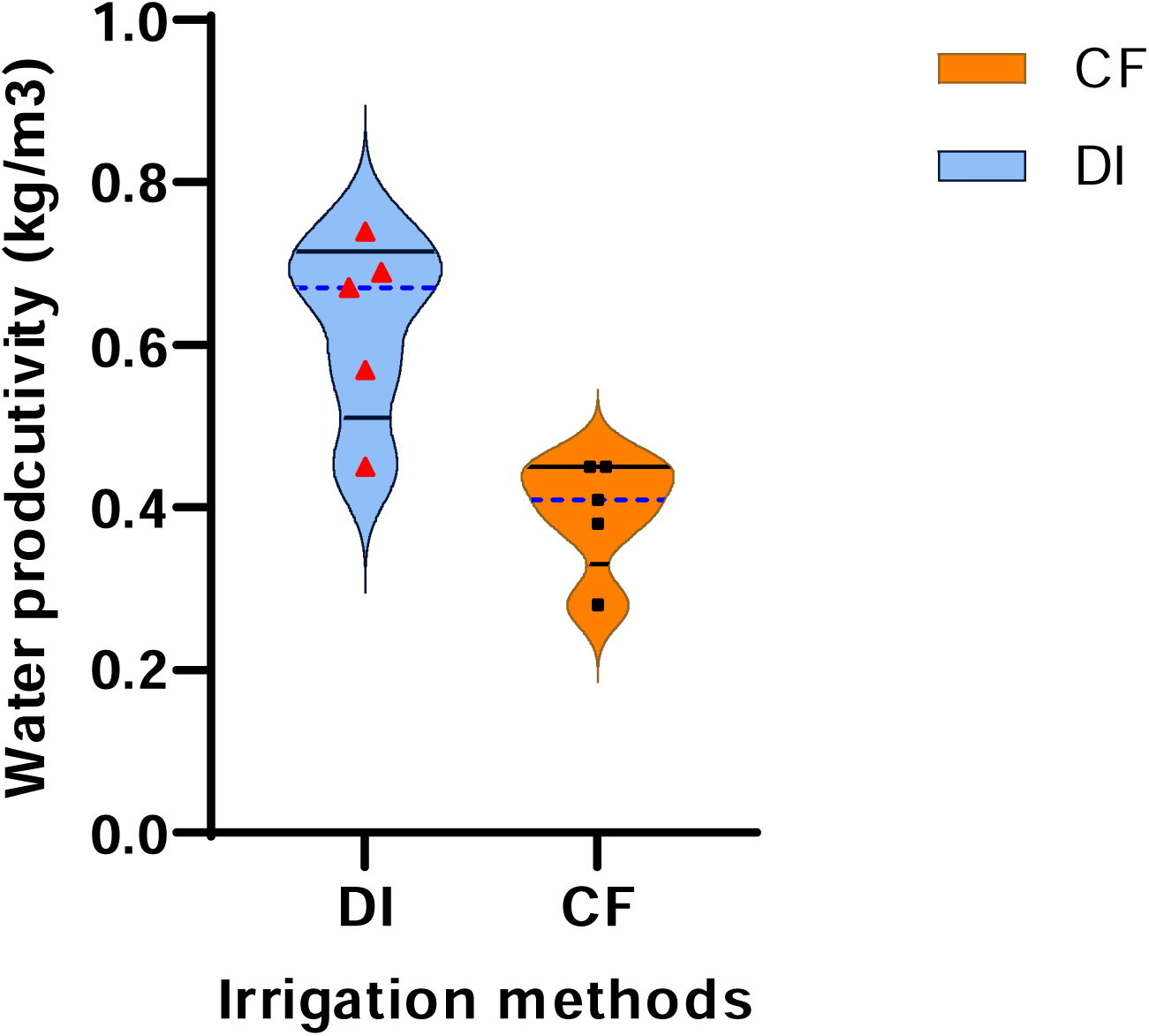
Water productivity (kg/m^3^) under continuous flooding (CF) and drip irrigation (DI) for five sites in Sathyamangalam, Tamil Nadu, India

### Methane emissions, global warming potential and yield scaled-global warming potential

Analysis of trial data indicated that methane emission (kg ha^-1^season^-1^), global warming potential (GWP) of methane (kg CO_2_ eq./ha) and yield-scaled global warming potential (kg CO_2_ eq./kg) were significantly influenced by irrigation methods, fertiliser management and varieties selection. Among the eight treatments, T5 exhibited the lowest emissions (CH_4_ - 91.9 kg/ha/ season, GWP – 2,574 kg CO_2_ eq./ha, YS-GWP - 0.5 kg.CO_2_ eq./kg), whereas the highest emissions were recorded in T4 (CH_4_ - 538.9 kg/ha/ season, GWP - 15,088 kg CO_2_ eq./ha, YS-GWP - 2.8 kg.CO_2_ eq./kg^-^) (Figure 6a-c). Drip irrigation significantly reduced methane emissions by over 60% compared to continuous flooding, with mean CH_4_: 128.7 versus 402.3 kg/ha/season, GWP: 3,603 versus 11,264.9 kg CO_2_ eq./ha, YS-GWP: 0.7 versus 2.1 kg.CO_2_ eq./kg. These differences were statistically significant across all sites (p < 0.01) (Supplemental Figure 1a-c).

**Figure 6:**
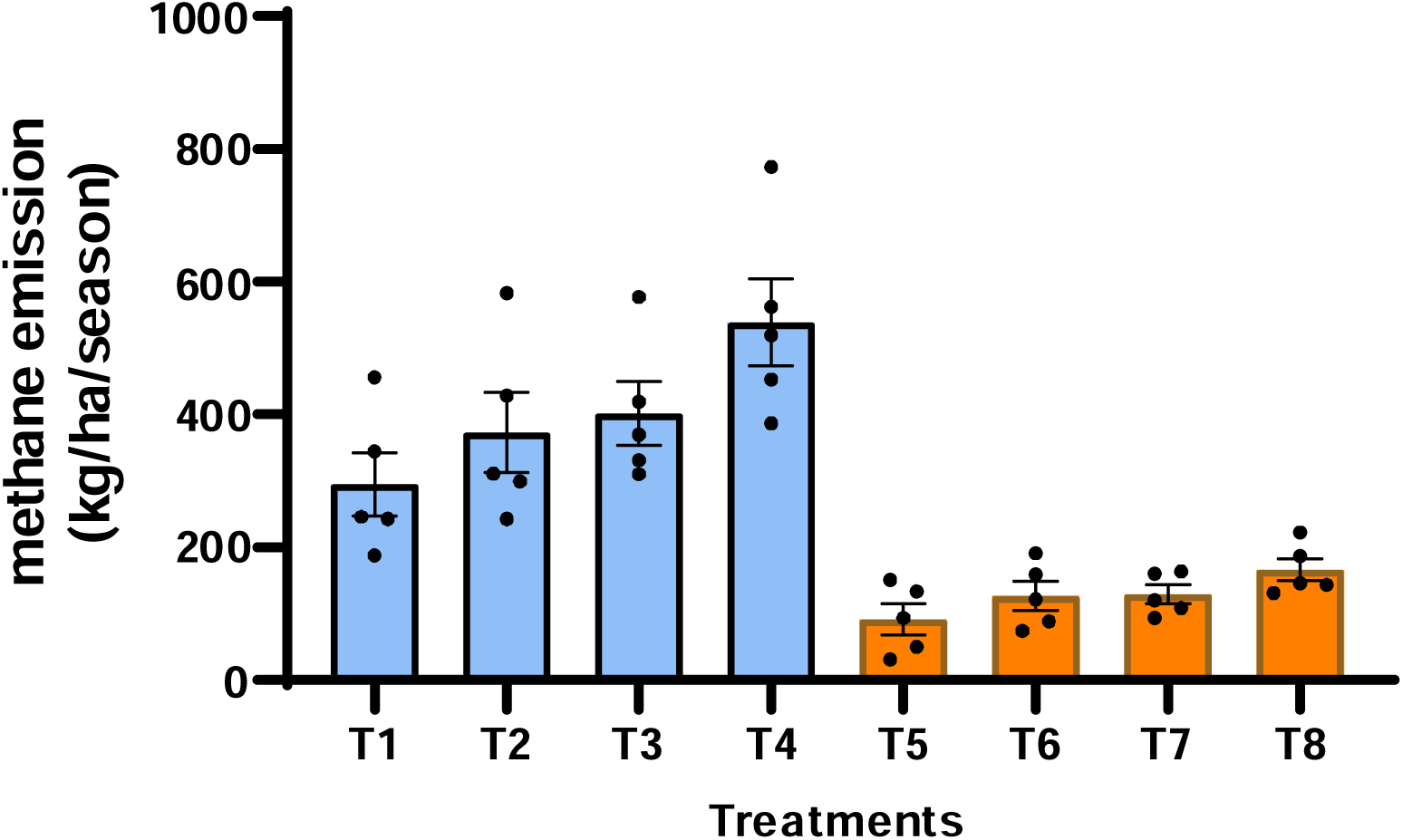
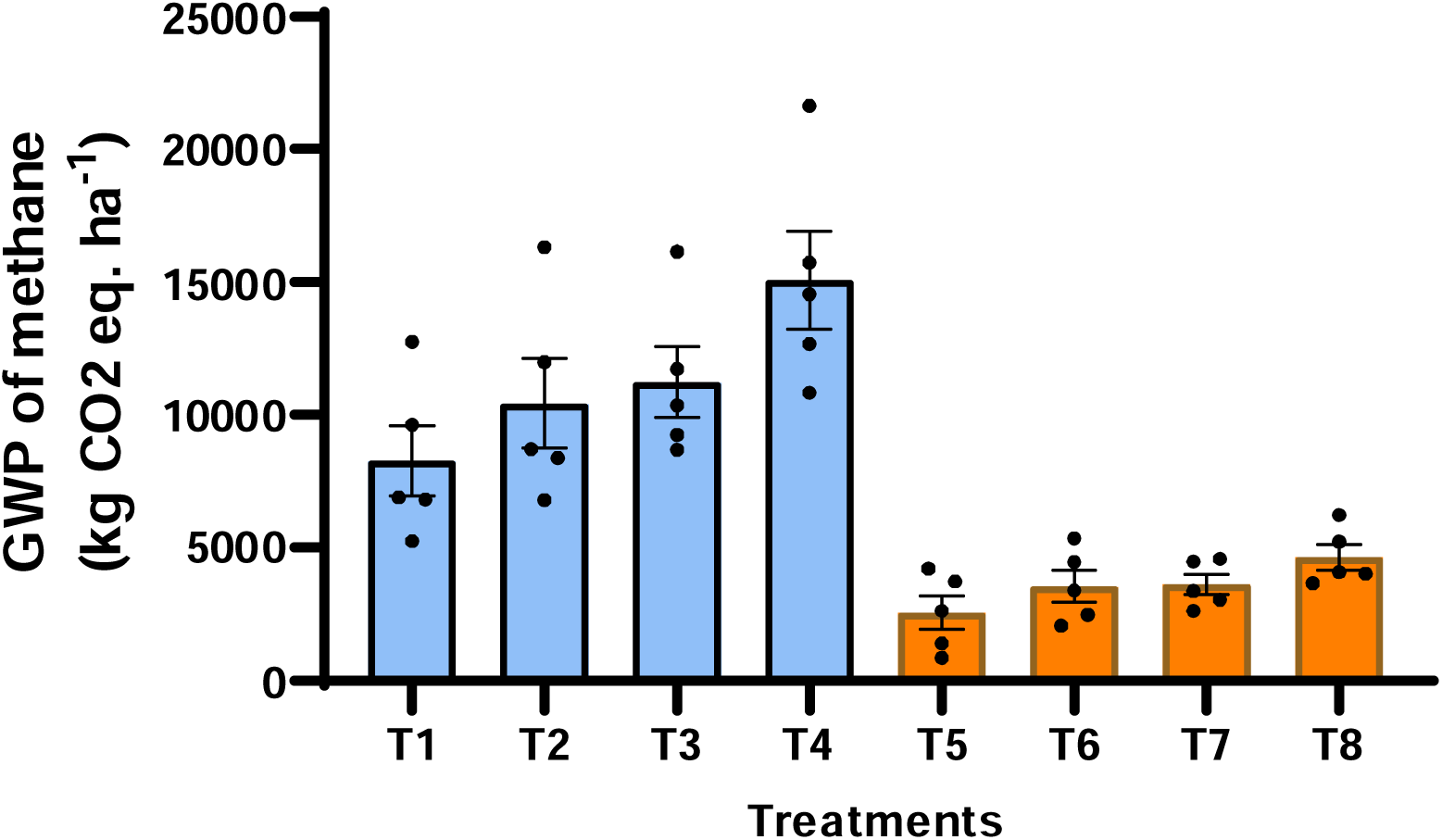
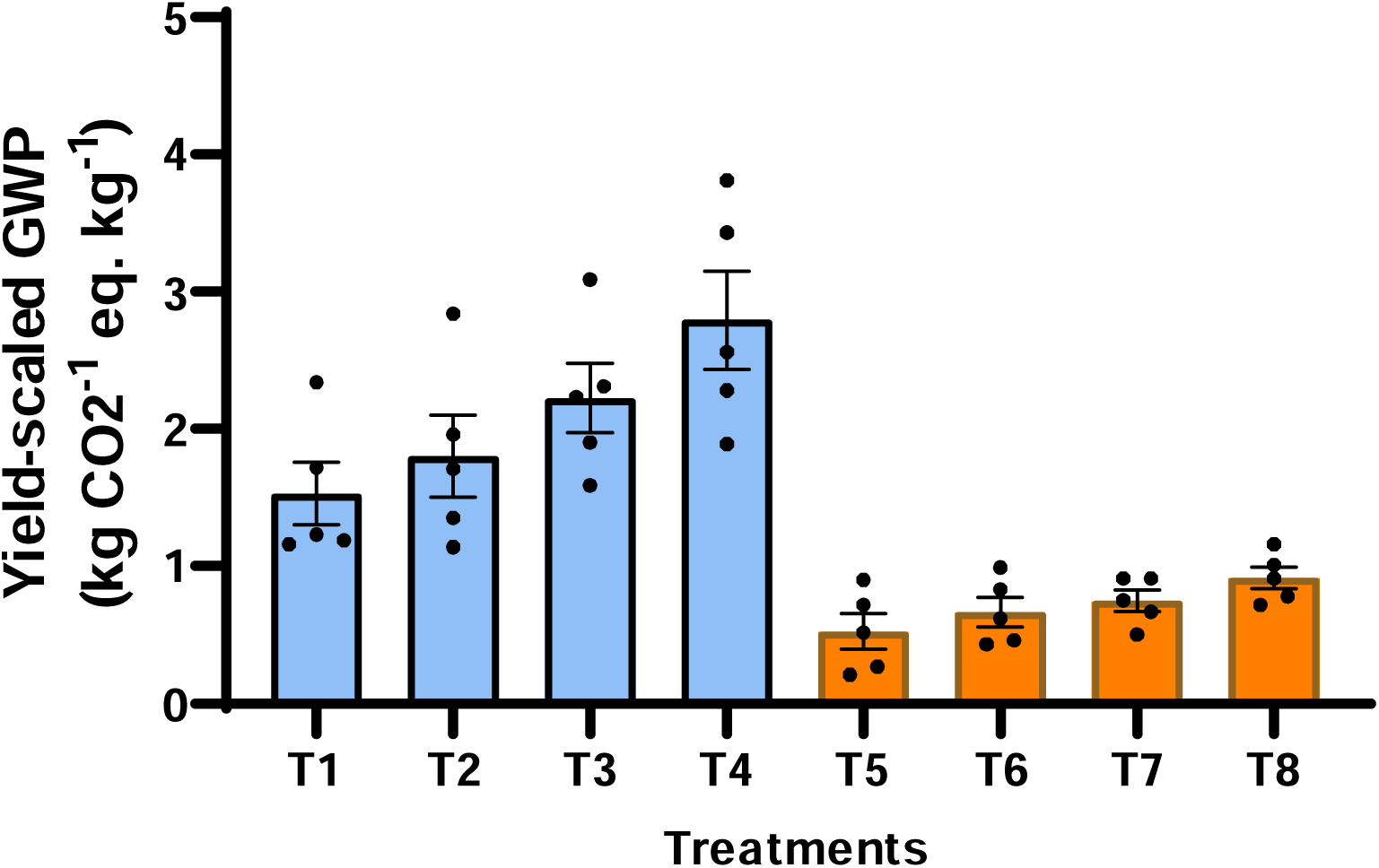
Average data for methane emission, global warming potential (GWP) and yield-scaled global warming potential for eight treatments carried out in five site in Sathyamangalam, Tamil Nadu, India. **(a)** Data represents average methane emission analysis across five sites, each subjected to eight distinct treatments. In the graph, each bar represents the mean methane emission for a given treatment, while “• “indicates the corresponding emission values for individual sites. Treatments T1 – T4 represent continuous flooding irrigation, incorporating two fertilizer types and two varieties, T5 – T8 is corresponds to drip irrigation, also with two fertiliser and two varieties. ANOVA analysis confirms a significant overall treatment effect at p value <0.001. **(b)** Data represents average global warming potential of methane across five sites, each subjected to eight distinct treatments. In the graph, each bar represents the mean global warming potential for a given treatment, while “• “indicates the corresponding emission values for individual sites. Treatments T1 – T4 represent continuous flooding irrigation, incorporating two fertilizer types and two varieties, T5 – T8 is corresponds to drip irrigation, also with two fertiliser and two varieties. ANOVA analysis confirms a significant overall treatment effect at p value <0.001. **(c)** Data represents average yield-scaled global warming potential values across five sites, each subjected to eight distinct treatments. In the graph, each bar represents the mean yield-scaled global warming potential for a given treatment, while “• “indicates the corresponding emission values for individual sites. Treatments T1 – T4 represent continuous flooding irrigation, incorporating two fertilizer types and two varieties, T5 – T8 is corresponds to drip irrigation, also with two fertiliser and two varieties. ANOVA analysis confirms a significant overall treatment effect at p value <0.001.

Fertiliser choice also significantly influenced the emissions, TLL-F outperforming conventional farmers’ practices (FP). Mean methane emissions were 221.9 versus 309.0 kg. /ha/season, GWP was 6,215 versus 8,653 kg CO_2_ eq./ha, YS-GWP was 1.1 vs 1.7 kg.CO_2_ eq./kg, likely due to enhanced nutrient synchrony and suppressed methanogenic activity under sulfate-based fertiliser combinations. The varietal differences were also evident: V1-ADT-45 emitted less (CH_4_: 229.7 kg. /ha/season, GWP: 6,430 kg CO_2_ eq./ha, YS-GWP: 1.3 kg.CO_2_ eq./kg) than BPT-5204 (CH_4_: 301 kg./ha/season, GWP: 8,438 kg CO_2_ eq./ha, YS-GWP: 1.5 kg.CO_2_ eq./kg), (Supplemental Table 4), potentially due to its shorter crop duration and lower root exudation.

The effects of fertiliser combination and rice varieties on methane emissions were highly site-specific, with complex interactions observed at sites 4 and 5. Treatment T4 (CF + FP + BPT-5204) consistently produced the highest methane emissions (773–520 kg./ha), whereas drip-irrigation based treatments (T5–T7) recorded the lowest emissions (31–151 kg./ha), underscoring the effectiveness of drip irrigation in limiting anaerobic conditions favourable for methanogenesis. Parthasarthi et. al. (2019) similarly reported a ∼ 49% reduction in yield-scaled global warming potential under drip irrigation compared to conventional continuous flooding. Under continuous flooding, oxygen-deprived microsites enable methanogenic archaea to convert soil carbon into methane, whereas drip irrigation systems maintain aerobic zones that suppress methanogens and promote oxidative pathways. Methane under drip irrigation and subsurface drip irrigation (SDI) have been attributed to increases methanotroph populations by approximately 39.8% (Parthasarathi et al., 2019, Jiao et al.,2006, Oo et al., 2018a).

The fertiliser combination TLL-F reduced methane emissions by 20–45% compared to conventional farmer practice, with treatment 5 (DI + TLL-F) achieving low emissions while maintaining stable yields, likely due to enhanced nutrient-use efficiency and improved root zone aeration. In contrast, T4 produced high yields but the cost of substantially higher greenhouse gas emissions (Figure 6). Sulfate -based amendments such as gypsum suppressed methane production by offering alternative electron acceptors (sulphate, Fe, Mn) as reported by Van Breemen & Feijtel (1990), Danier van der Gon & Neue (1994), Ro et al. (2011), Theint et al. (2016), Kowshika et al. (2019), and Jun-Hong et al. (2016), the latter observing a 60% decline in methane flux. Specifically, Danier van der Gon et al. (1994) demonstrated that gypsum applications at 6.66 Mg/Ha reduced methane emissions by 54% - 71% in paddy field. Varietal effects were notable, with BPT-5204 emitting approximately 100 kg CH_4_ /ha^-1^ /season^-1^ more than ADT-45, attributable to its longer growth duration, higher biomass, and greater carbon exudation. Slamento et al. (2024) reported that short-duration rice varieties emit 25-30% less methane than longer-duration varieties, due to reduced cumulative exposure and lower total organic matte available for methanogens. These observations are consistent with earlier studies, linking elevated methane emissions and global warming potential to longer cultivation period and high biomass cultivar (Wang et al. 2021, Gogoi et al. 2008; Baruah et al. 2010; Khosa et al. 2010; Bharali et al. 2017; Bhattacharyya et al. 2019). Overall, genotypes, fertiliser regime, and water management interact synergistically to regulate methane fluxes in rice cultivation systems.

### Conclusions

This integrated analysis underscores the critical role of irrigation method, nutrient management, and varietal selection in enhancing rice productivity while mitigating methane emissions across diverse agro-ecological contexts. Drip irrigation consistently outperformed continuous flooding, not only reducing methane emissions but also improving the water use efficiency and crop yields which are key components of food security. The combination of drip irrigation with optimised fertiliser management (DI+TLL-F) and climate-smart varieties such as ADT-45 significantly reduces both absolute methane flux and yield-scaled global warming potential (YS-GWP), supporting the shift toward sustainable intensification. In contrast, traditional systems particularly T4 (CF+FP+BPT-5204), resulted in disproportionately high methane emissions due to greater biomass accumulation, prolonged anaerobic periods, and inefficient nutrient utilization. The observed site-specific variability in emission dynamics highlights the importance of tailoring management and practices to local agro-climatic, and varietal conditions. Future research should integrate dynamic soil parameters, evapotranspiration patterns, and crop physiological responses to refine emission, estimates and develop stage-specific mitigation strategies. Collectively, these findings advocate for a systematic approach to rice cultivation, that combines precision irrigation, balanced nutrition management, and careful varietal evaluation to enhance productivity while aligning with environmentally sustainable goals, thereby contributing meaningfully to both national and global climate objectives.

## Acknowledgements

We acknowledge research grants from Temasek Foundation, Singapore and the Temasek Life Sciences Laboratory, Singapore. We also would like to thank Dr Goh Phuay Yee for her invaluable support in project management and for the comments and suggestions for this manuscript.

## Contribution of authors

K.V – Field trial design and data analysis; original draft preparation; A. M, M.S & K.L– experiment designs and sample analysis; K.S – original draft preparation and data analysis; ZY & N.N – original draft and scientific inputs for trials, S.R – Field trial design, supervisor, and original draft preparation and review. All the authors have read the manuscript and agreed for publication.

## Supplemental Data

### Supplemental Materials

**Table 1:** Rice plant growth, yield and methane data averaged across five sites in Sathyamangalam.

| Treatment | Plant height (cm) | Effective tillers/plant | Panicle length (cm) | Grain/panicle | Sterile spikelet (numbers) | Unfilled grains (%) | Test weight (1,000 seeds) | Grain yield (tonnes/ha) | Straw yield (in tonnes/ha) | Harvest index |
| --- | --- | --- | --- | --- | --- | --- | --- | --- | --- | --- |
| Irrigation |  |  |  |  |  |  |  |  |  |  |
| CF (I1) | 93.56 | 5.57 | 20.08 | 158.50 | 18.62 | 10.67 | 15.64 | 5.46 | 5.02 | 0.50 |
| DI (I2) | 91.61 | 6.95 | 20.23 | 149.17 | 23.41 | 13.99 | 14.99 | 5.02 | 4.76 | 0.48 |
| Fertilisers |  |  |  |  |  |  |  |  |  |  |
| TLL-RDF (F1) | 92.18 | 7.16 | 19.99 | 153.90 | 17.59 | 11.37 | 15.39 | 5.35 | 4.92 | 0.49 |
| FP (F2) | 92.99 | 7.36 | 20.32 | 153.77 | 24.44 | 13.29 | 15.24 | 5.14 | 4.87 | 0.48 |
| Varieties |  |  |  |  |  |  |  |  |  |  |
| ADT-45 (V1) | 91.95 | 6.80 | 19.63 | 162.40 | 19.58 | 11.80 | 16.52 | 5.05 | 4.34 | 0.47 |
| BPT-5204 (V2) | 93.22 | 7.72 | 20.68 | 145.27 | 22.46 | 12.86 | 14.11 | 5.43 | 5.45 | 0.50 |
| *Irrigation (I)* | * | ** | ns | * | ** | ** | ** | ** | ns | ** |
| *Fertilisers (F)* | ns | ns | * | ns | ** | ** | * | ** | ns | ** |
| *Varieties (V)* | ns | ** | ** | ** | ** | * | ** | ** | ** | ** |
| *V x I* | ns | ns | ns | ns | ns | ns | ** | ns | ns | ns |
| *V x F x I* | ns | ns | ns | ns | ns | ns | * | ns | ns | Ns |
| *F x I* | ns | ns | ns | ns | ns | ns | ns | * | ns | ns |

*\*- Significant at 5%, \*\*- Significant at 1%, ns-nonsignificant. In a column, figures having the same letter(s) do not differ significantly, whereas figures with different letter(s) differ significantly, as per LSD at 5% level of significance.*

**Supplemental Table 2:**
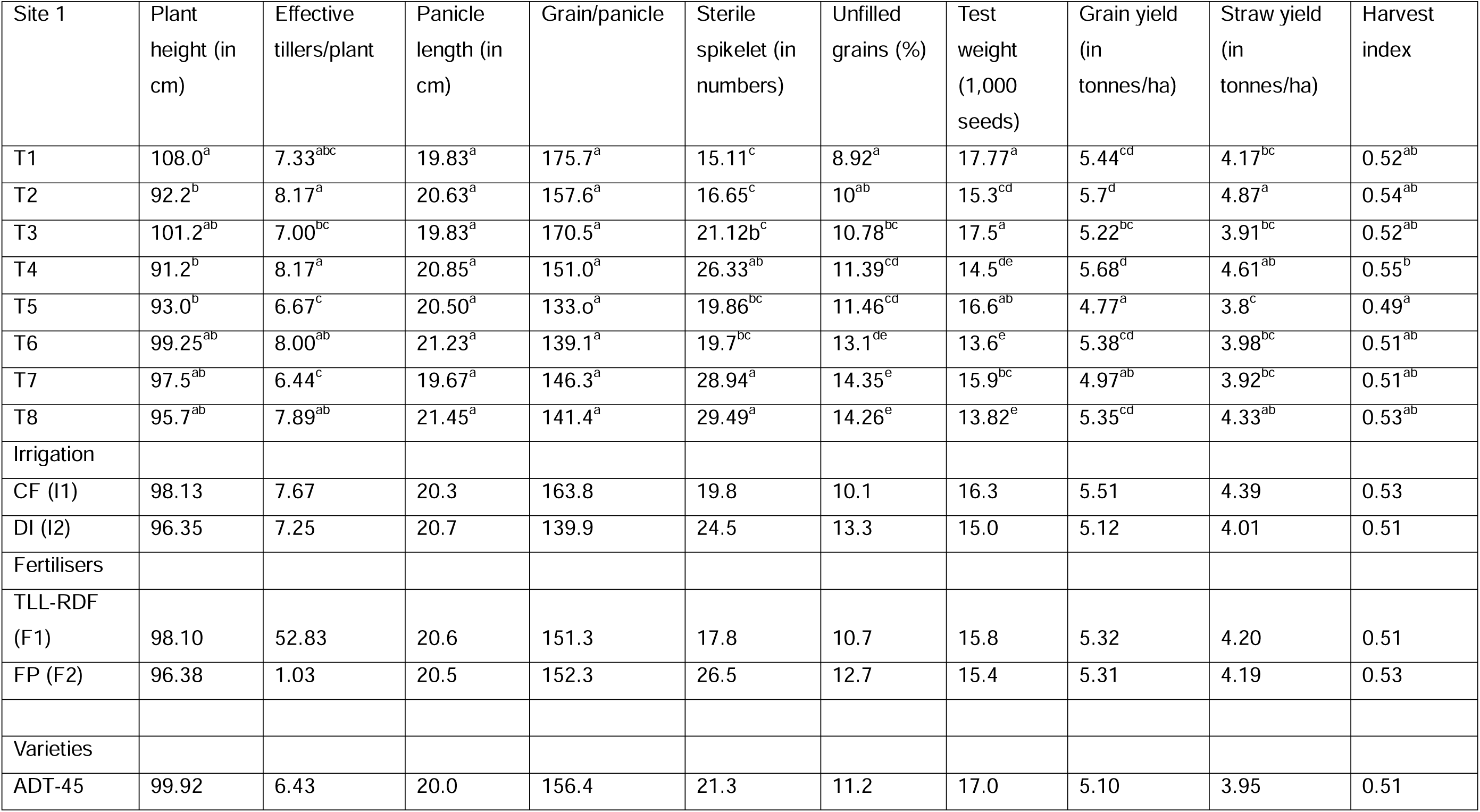
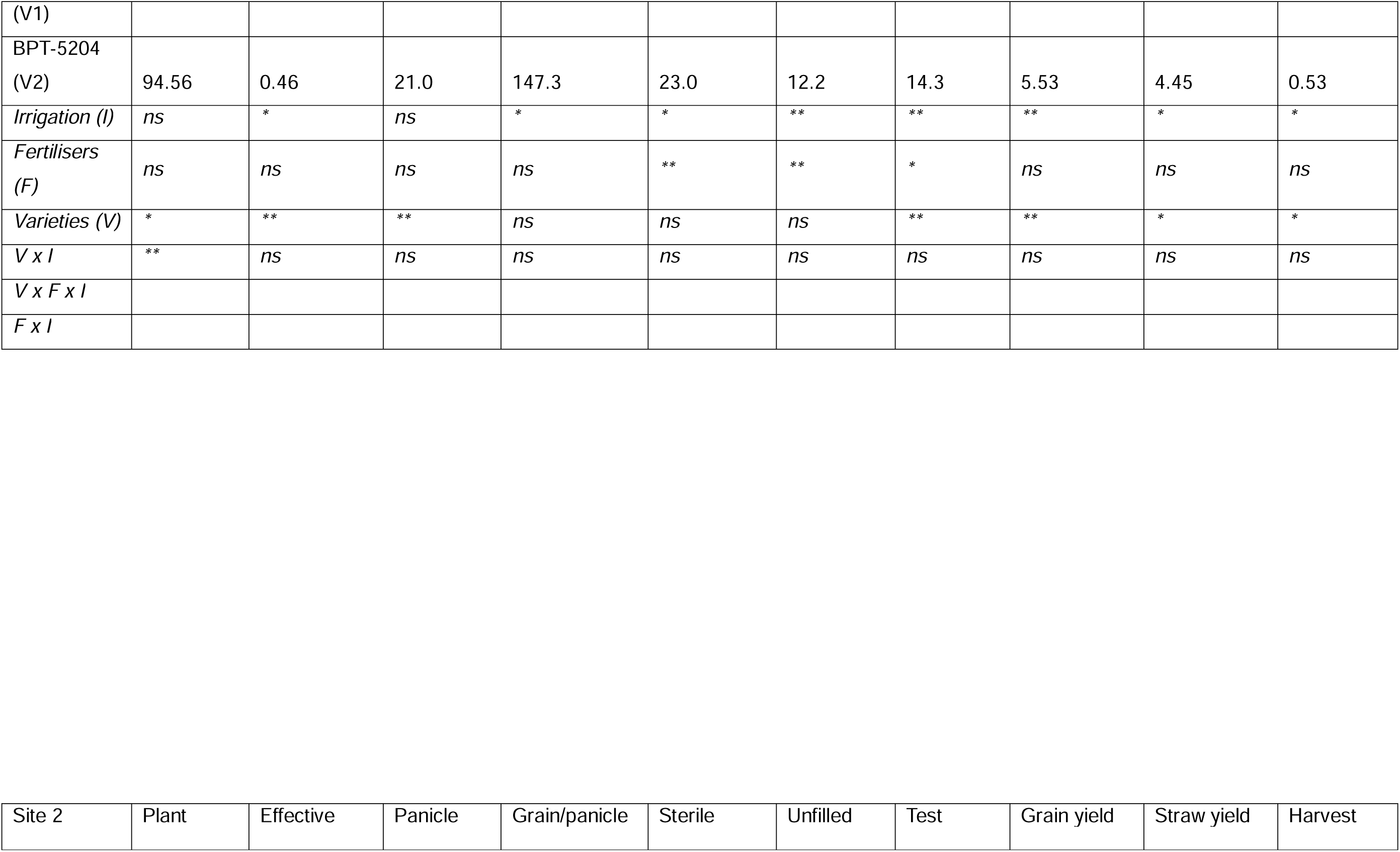
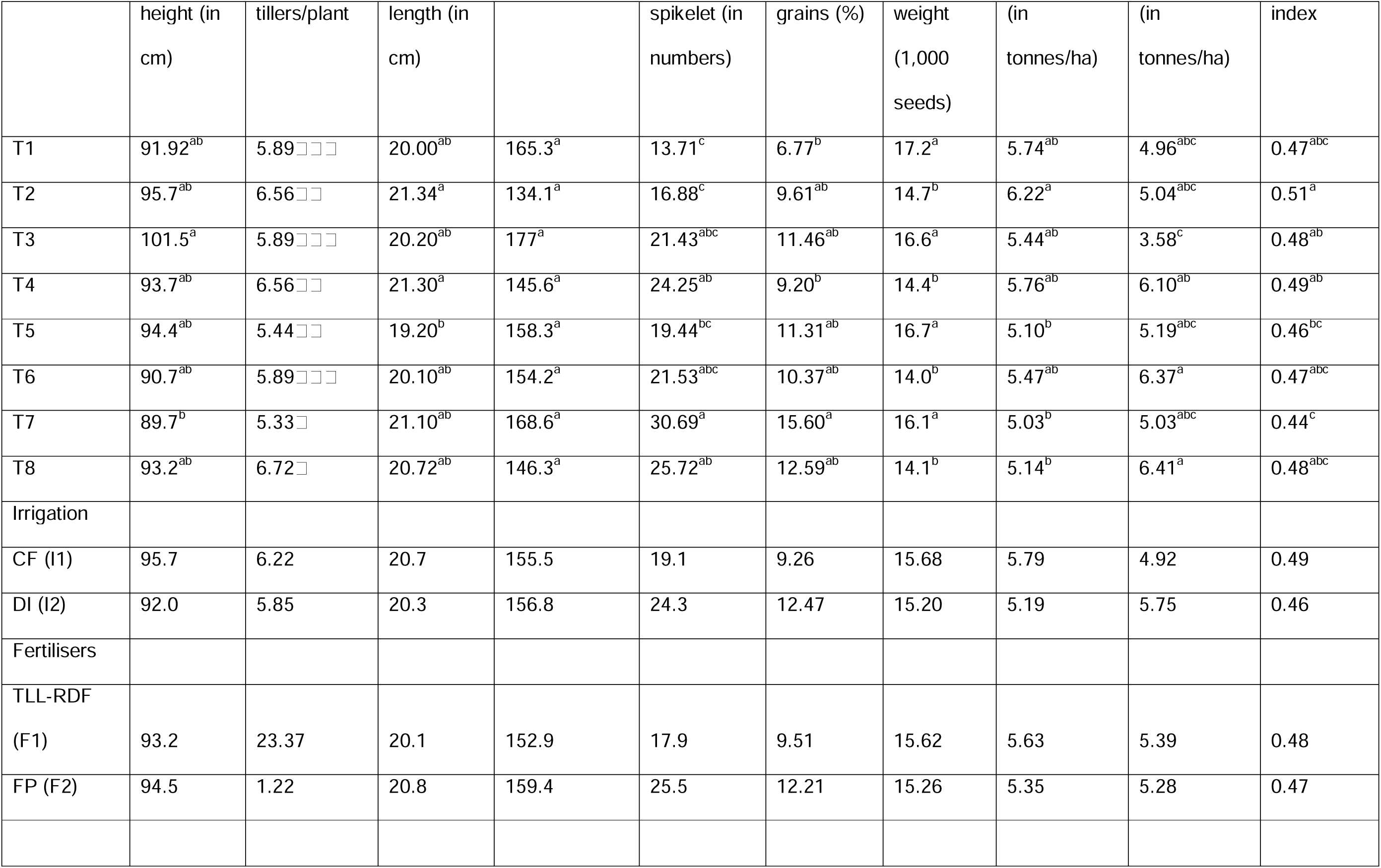
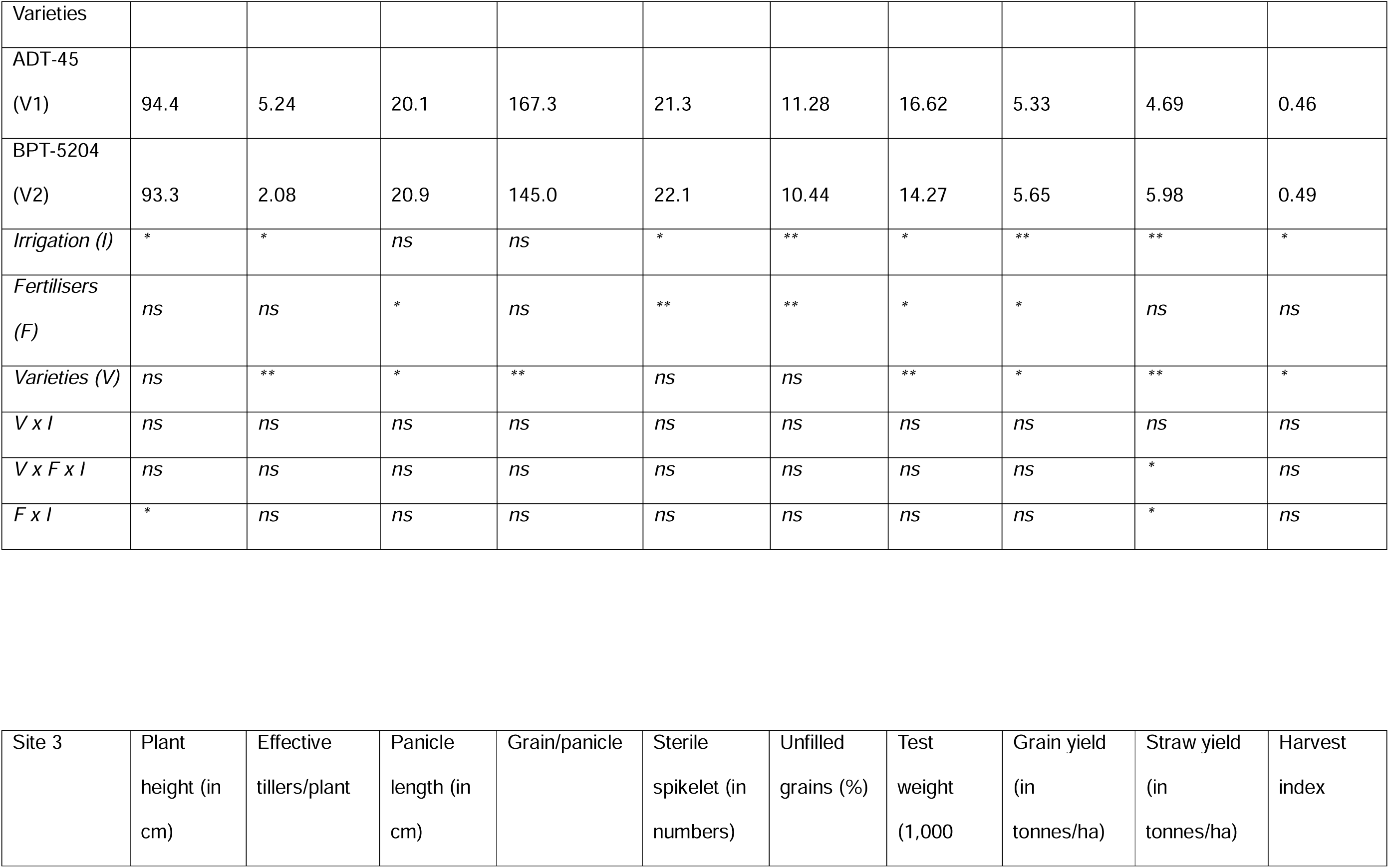
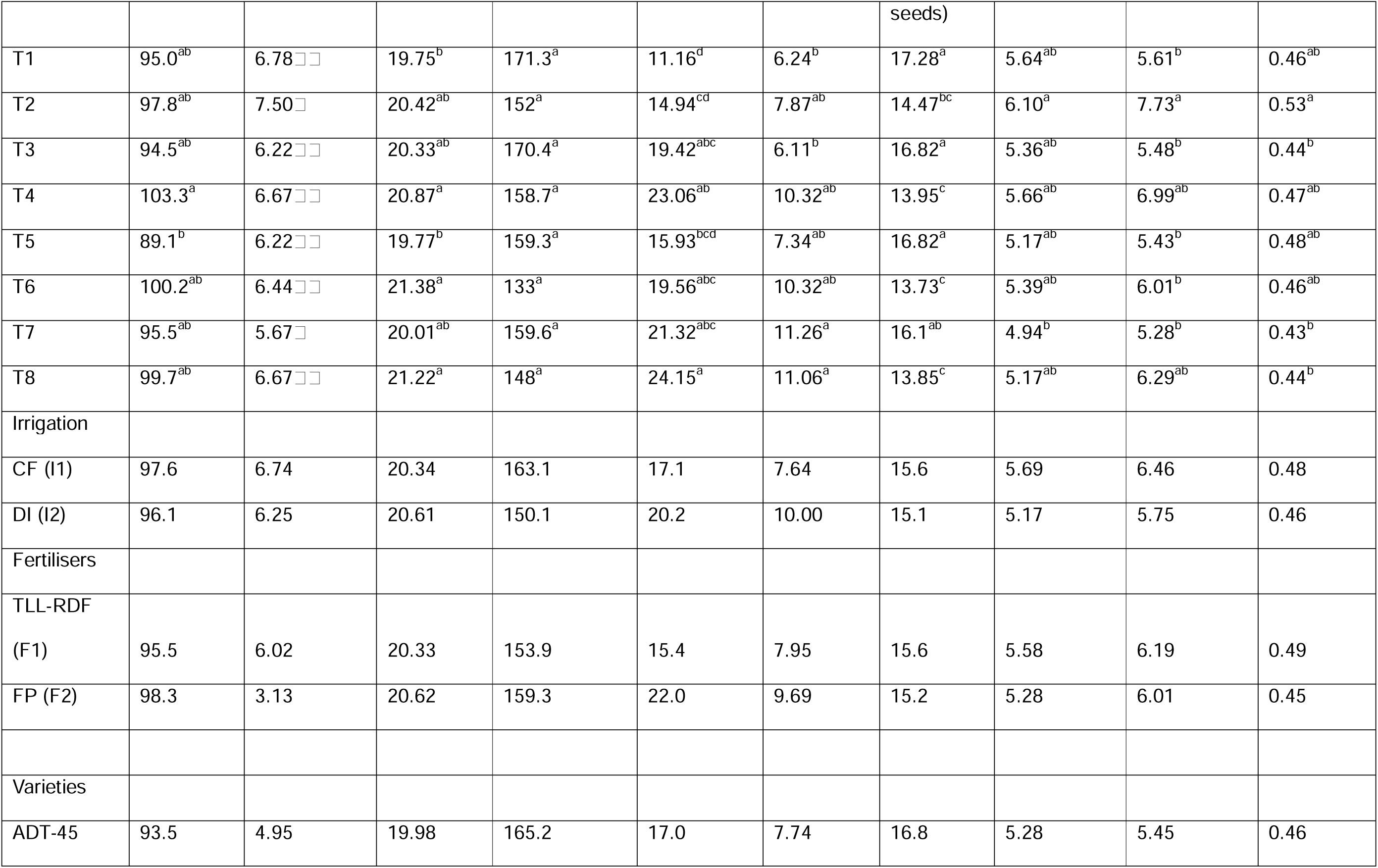
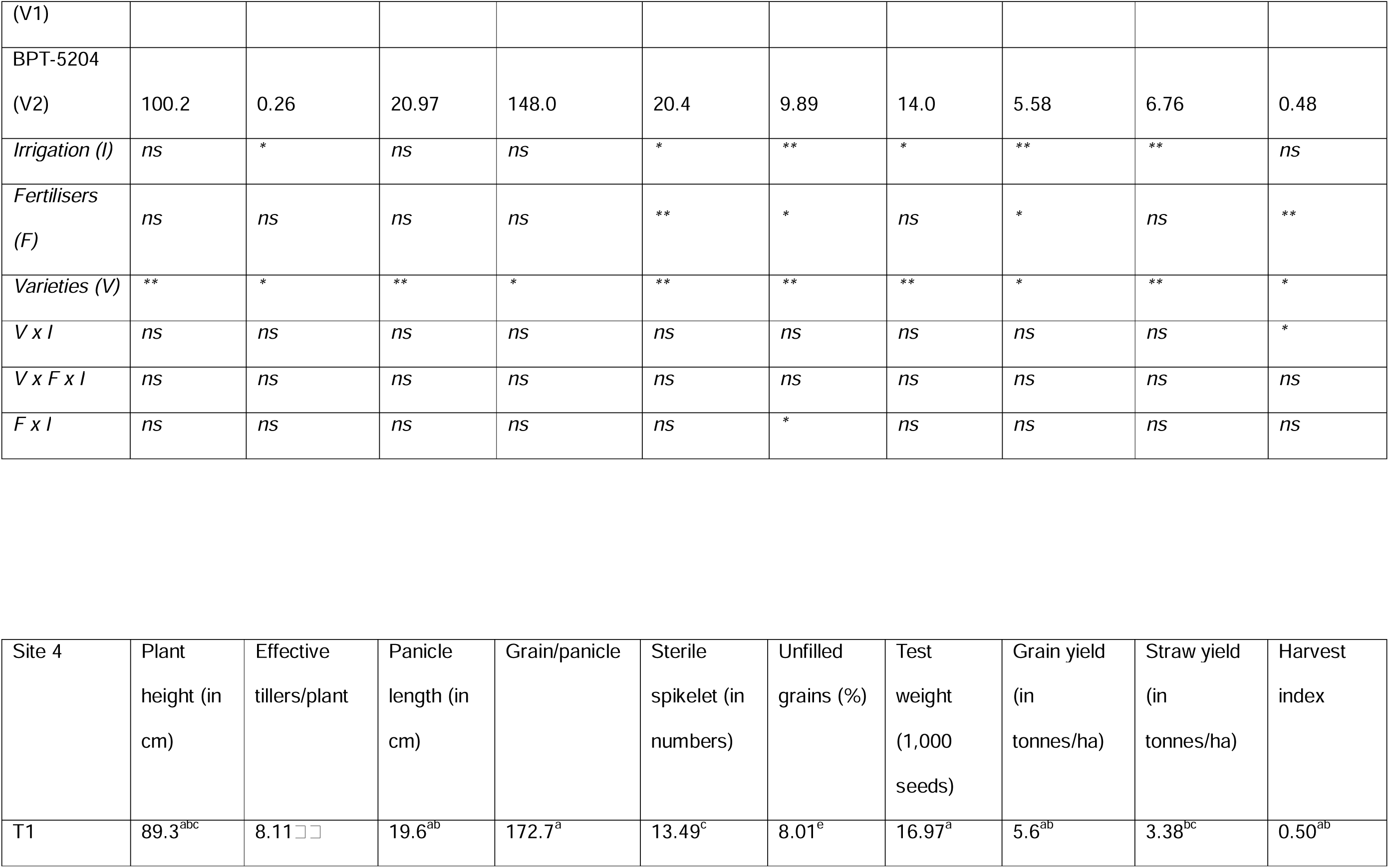
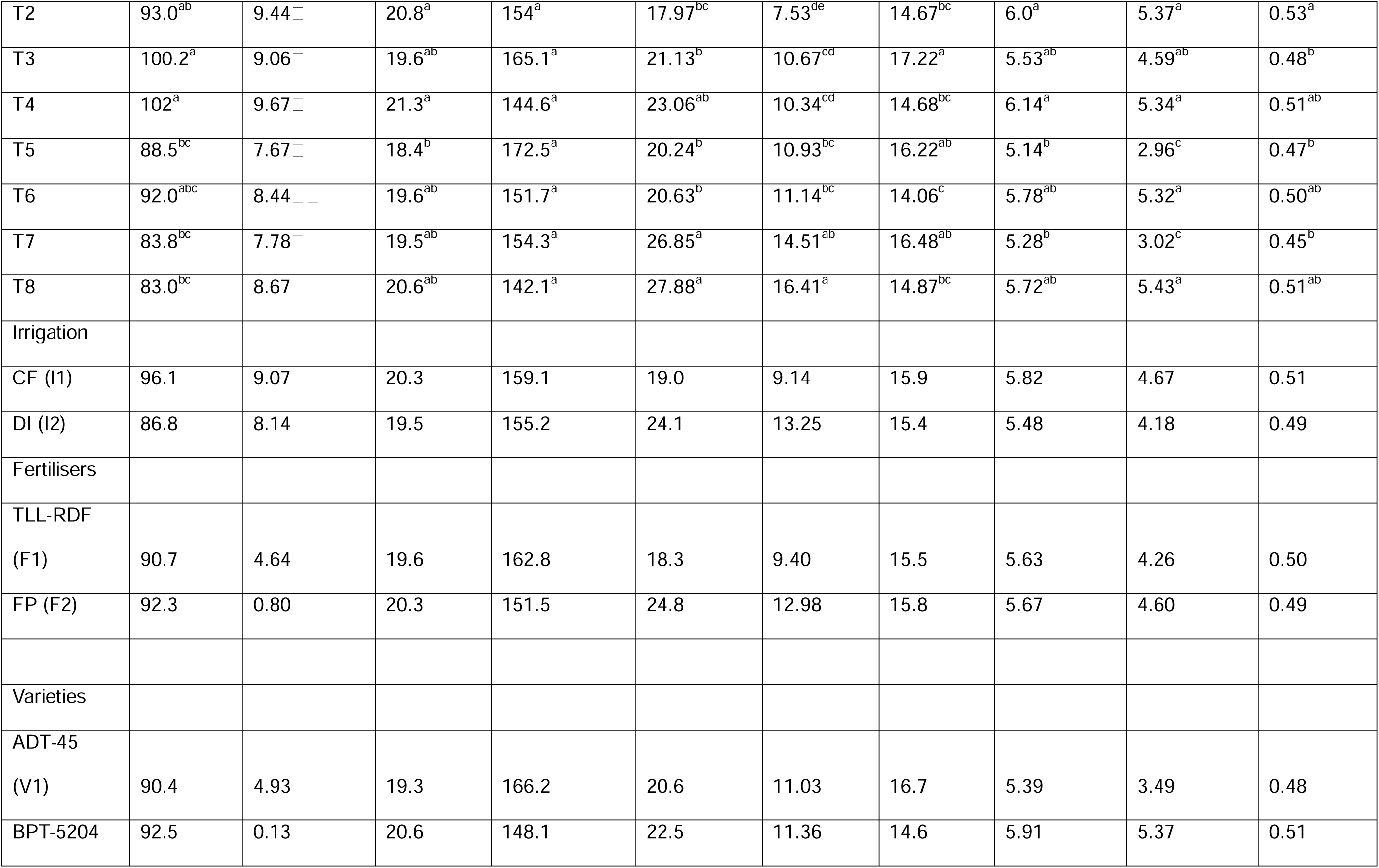
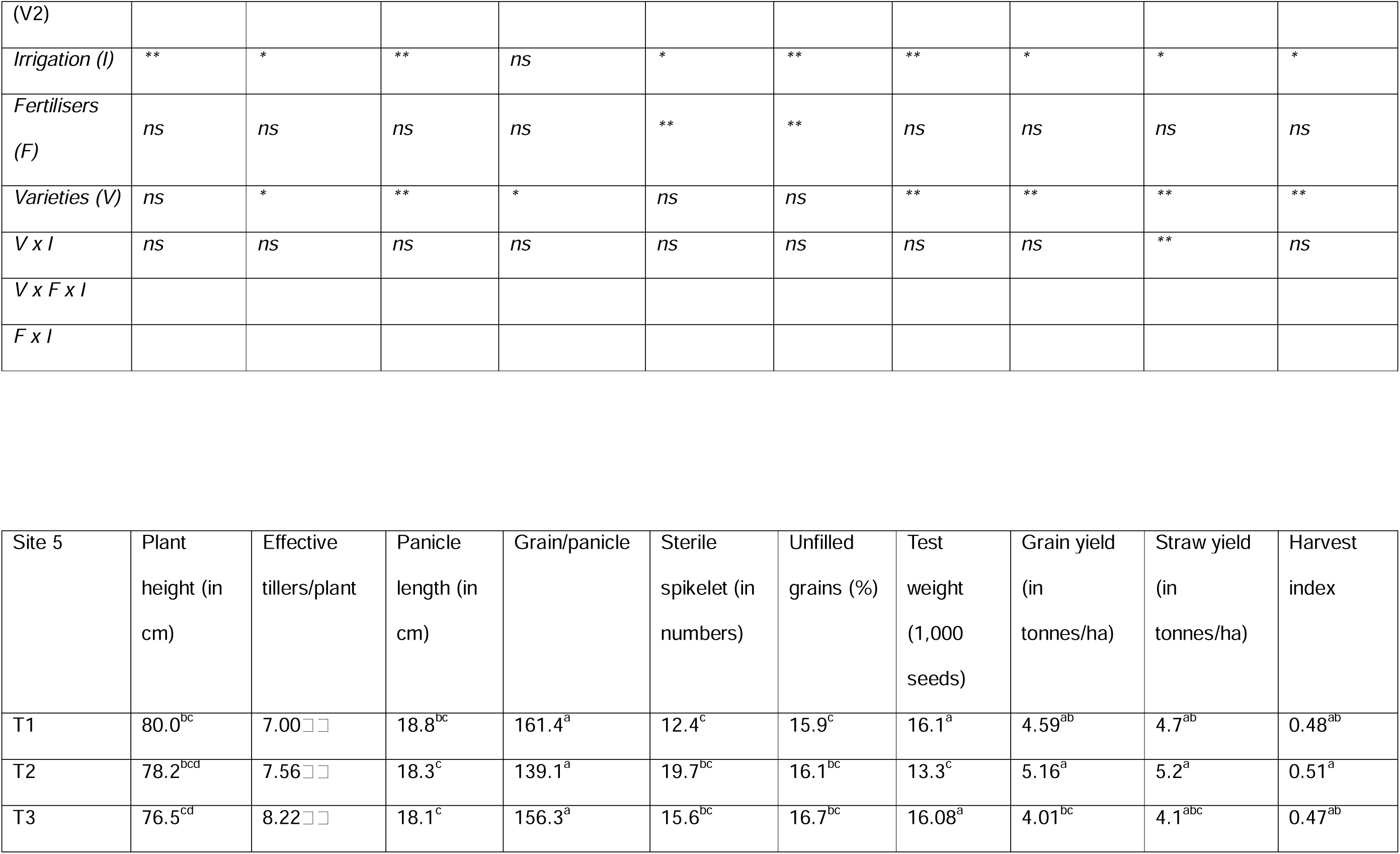
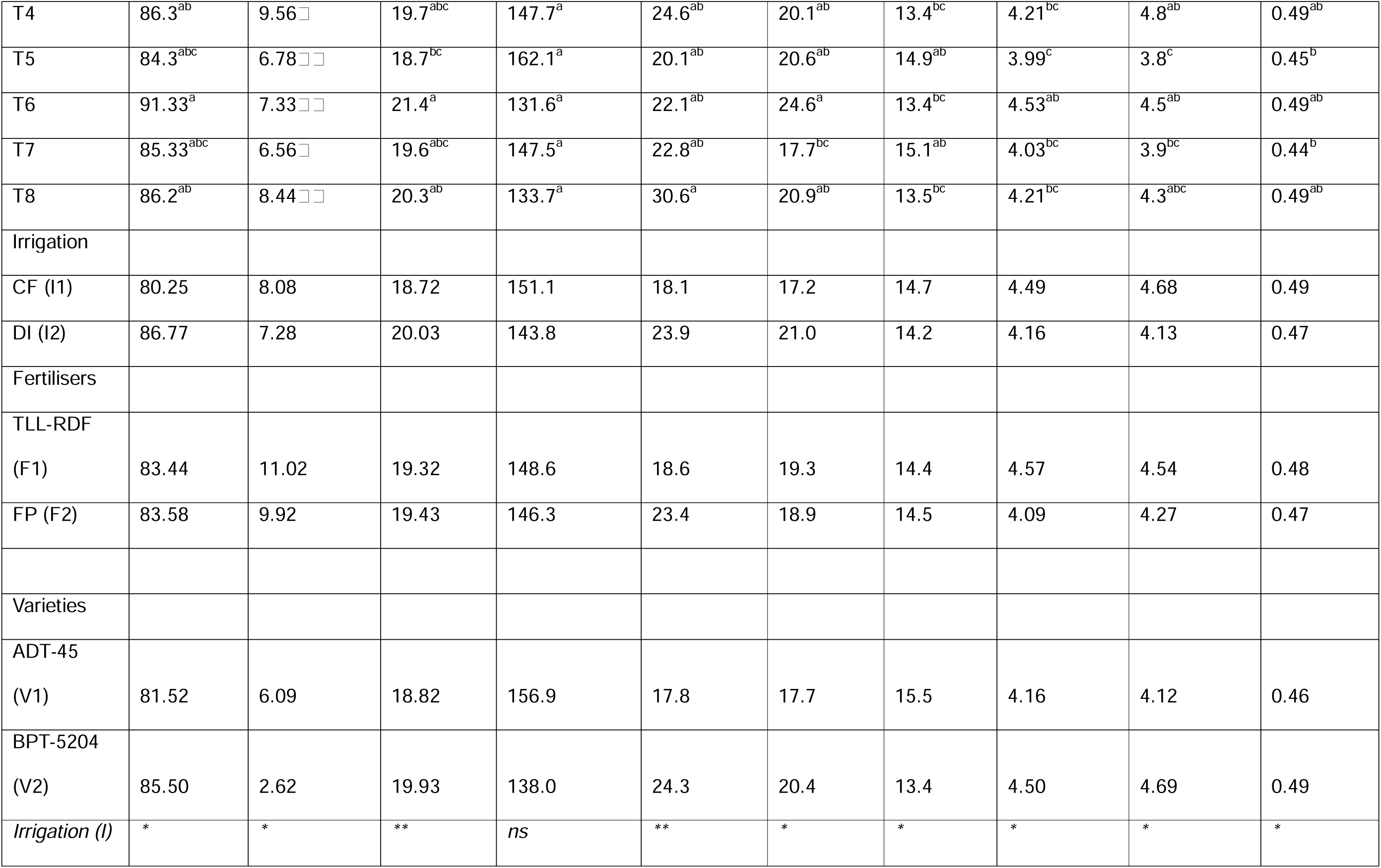
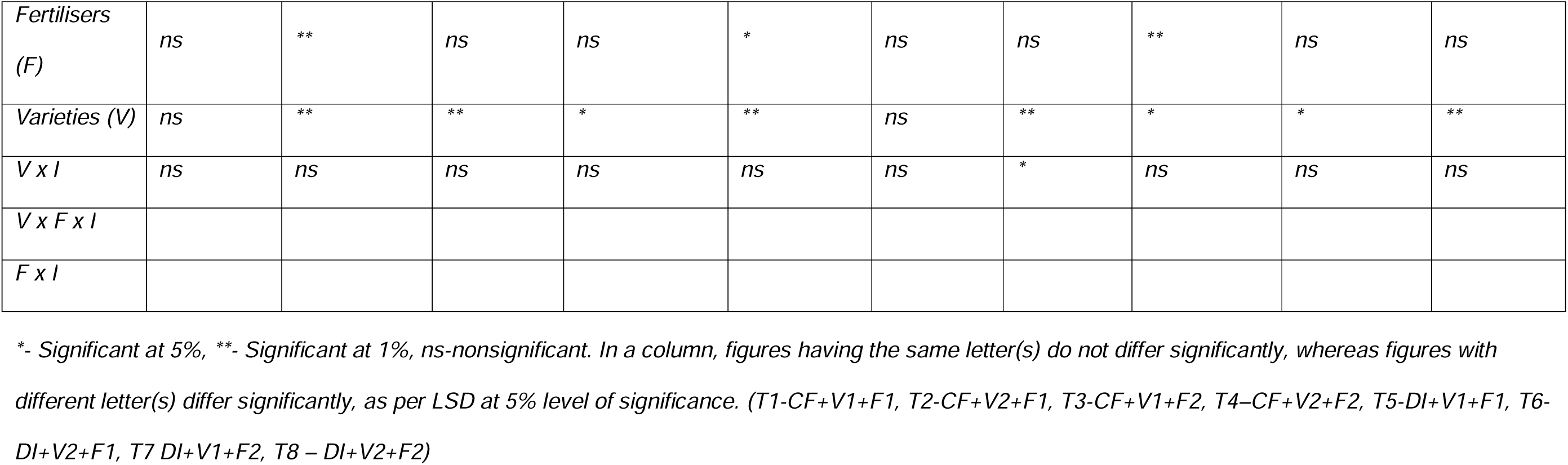
Site-wise data of plant and yield attributes in rice varieties under 2 irrigation systems and 2 fertiliser combinations (Site 1 to 5)

| Site 1 | Plant height (in cm) | Effective tillers/plant | Panicle length (in cm) | Grain/panicle | Sterile spikelet (in numbers) | Unfilled grains (%) | Test weight (1,000 seeds) | Grain yield (in tonnes/ha) | Straw yield (in tonnes/ha) | Harvest index |
| --- | --- | --- | --- | --- | --- | --- | --- | --- | --- | --- |
| T1 | 108.0<sup>a</sup> | 7.33<sup>abc</sup> | 19.83<sup>a</sup> | 175.7<sup>a</sup> | 15.11<sup>c</sup> | 8.92<sup>a</sup> | 17.77<sup>a</sup> | 5.44<sup>cd</sup> | 4.17<sup>bc</sup> | 0.52<sup>ab</sup> |
| T2 | 92.2<sup>b</sup> | 8.17<sup>a</sup> | 20.63<sup>a</sup> | 157.6<sup>a</sup> | 16.65<sup>c</sup> | 10<sup>ab</sup> | 15.3<sup>cd</sup> | 5.7<sup>d</sup> | 4.87<sup>a</sup> | 0.54<sup>ab</sup> |
| T3 | 101.2<sup>ab</sup> | 7.00<sup>bc</sup> | 19.83<sup>a</sup> | 170.5<sup>a</sup> | 21.12<sup>b</sup><sup>c</sup> | 10.78<sup>bc</sup> | 17.5<sup>a</sup> | 5.22<sup>bc</sup> | 3.91<sup>bc</sup> | 0.52<sup>ab</sup> |
| T4 | 91.2<sup>b</sup> | 8.17<sup>a</sup> | 20.85<sup>a</sup> | 151.0<sup>a</sup> | 26.33<sup>ab</sup> | 11.39<sup>cd</sup> | 14.5<sup>de</sup> | 5.68<sup>d</sup> | 4.61<sup>ab</sup> | 0.55<sup>b</sup> |
| T5 | 93.0<sup>b</sup> | 6.67<sup>c</sup> | 20.50<sup>a</sup> | 133.0<sup>a</sup> | 19.86<sup>bc</sup> | 11.46<sup>cd</sup> | 16.6<sup>ab</sup> | 4.77<sup>a</sup> | 3.8<sup>c</sup> | 0.49<sup>a</sup> |
| T6 | 99.25<sup>ab</sup> | 8.00<sup>ab</sup> | 21.23<sup>a</sup> | 139.1<sup>a</sup> | 19.7<sup>bc</sup> | 13.1<sup>de</sup> | 13.6<sup>e</sup> | 5.38<sup>cd</sup> | 3.98<sup>bc</sup> | 0.51<sup>ab</sup> |
| T7 | 97.5<sup>ab</sup> | 6.44<sup>c</sup> | 19.67<sup>a</sup> | 146.3<sup>a</sup> | 28.94<sup>a</sup> | 14.35<sup>e</sup> | 15.9<sup>bc</sup> | 4.97<sup>ab</sup> | 3.92<sup>bc</sup> | 0.51<sup>ab</sup> |
| T8 | 95.7<sup>ab</sup> | 7.89<sup>ab</sup> | 21.45<sup>a</sup> | 141.4<sup>a</sup> | 29.49<sup>a</sup> | 14.26<sup>e</sup> | 13.82<sup>e</sup> | 5.35<sup>cd</sup> | 4.33<sup>ab</sup> | 0.53<sup>ab</sup> |
| Irrigation |  |  |  |  |  |  |  |  |  |  |
| CF (I1) | 98.13 | 7.67 | 20.3 | 163.8 | 19.8 | 10.1 | 16.3 | 5.51 | 4.39 | 0.53 |
| DI (I2) | 96.35 | 7.25 | 20.7 | 139.9 | 24.5 | 13.3 | 15.0 | 5.12 | 4.01 | 0.51 |
| Fertilisers |  |  |  |  |  |  |  |  |  |  |
| TLL-RDF (F1) | 98.10 | 52.83 | 20.6 | 151.3 | 17.8 | 10.7 | 15.8 | 5.32 | 4.20 | 0.51 |
| FP (F2) | 96.38 | 1.03 | 20.5 | 152.3 | 26.5 | 12.7 | 15.4 | 5.31 | 4.19 | 0.53 |
| Varieties |  |  |  |  |  |  |  |  |  |  |
| ADT-45 | 99.92 | 6.43 | 20.0 | 156.4 | 21.3 | 11.2 | 17.0 | 5.10 | 3.95 | 0.51 |
| (V1) |  |  |  |  |  |  |  |  |  |  |
| BPT-5204<br>(V2) | 94.56 | 0.46 | 21.0 | 147.3 | 23.0 | 12.2 | 14.3 | 5.53 | 4.45 | 0.53 |
| *Irrigation (I)* | *ns* | * | *ns* | * | * | ** | ** | ** | * | * |
| *Fertilisers (F)* | *ns* | *ns* | *ns* | *ns* | ** | ** | * | *ns* | *ns* | *ns* |
| *Varieties (V)* | * | ** | ** | *ns* | *ns* | *ns* | ** | ** | * | * |
| *V x I* | ** | *ns* | *ns* | *ns* | *ns* | *ns* | *ns* | *ns* | *ns* | *ns* |
| *V x F x I* |  |  |  |  |  |  |  |  |  |  |
| *F x I* |  |  |  |  |  |  |  |  |  |  |
| Site 2 | Plant | Effective | Panicle | Grain/panicle | Sterile | Unfilled | Test | Grain yield | Straw yield | Harvest |

|  | height (in<br>cm) | tillers/plant | length (in<br>cm) |  | spikelet (in<br>numbers) | grains (%) | weight<br>(1,000<br>seeds) | (in<br>tonnes/ha) | (in<br>tonnes/ha) | index |
| --- | --- | --- | --- | --- | --- | --- | --- | --- | --- | --- |
| T1 | 91.92<sup>ab</sup> | 5.89□□□ | 20.00<sup>ab</sup> | 165.3<sup>a</sup> | 13.71<sup>c</sup> | 6.77<sup>b</sup> | 17.2<sup>a</sup> | 5.74<sup>ab</sup> | 4.96<sup>abc</sup> | 0.47<sup>abc</sup> |
| T2 | 95.7<sup>ab</sup> | 6.56□□ | 21.34<sup>a</sup> | 134.1<sup>a</sup> | 16.88<sup>c</sup> | 9.61<sup>ab</sup> | 14.7<sup>b</sup> | 6.22<sup>a</sup> | 5.04<sup>abc</sup> | 0.51<sup>a</sup> |
| T3 | 101.5<sup>a</sup> | 5.89□□□ | 20.20<sup>ab</sup> | 177<sup>a</sup> | 21.43<sup>abc</sup> | 11.46<sup>ab</sup> | 16.6<sup>a</sup> | 5.44<sup>ab</sup> | 3.58<sup>c</sup> | 0.48<sup>ab</sup> |
| T4 | 93.7<sup>ab</sup> | 6.56□□ | 21.30<sup>a</sup> | 145.6<sup>a</sup> | 24.25<sup>ab</sup> | 9.20<sup>b</sup> | 14.4<sup>b</sup> | 5.76<sup>ab</sup> | 6.10<sup>ab</sup> | 0.49<sup>ab</sup> |
| T5 | 94.4<sup>ab</sup> | 5.44□□ | 19.20<sup>b</sup> | 158.3<sup>a</sup> | 19.44<sup>bc</sup> | 11.31<sup>ab</sup> | 16.7<sup>a</sup> | 5.10<sup>b</sup> | 5.19<sup>abc</sup> | 0.46<sup>bc</sup> |
| T6 | 90.7<sup>ab</sup> | 5.89□□□ | 20.10<sup>ab</sup> | 154.2<sup>a</sup> | 21.53<sup>abc</sup> | 10.37<sup>ab</sup> | 14.0<sup>b</sup> | 5.47<sup>ab</sup> | 6.37<sup>a</sup> | 0.47<sup>abc</sup> |
| T7 | 89.7<sup>b</sup> | 5.33□ | 21.10<sup>ab</sup> | 168.6<sup>a</sup> | 30.69<sup>a</sup> | 15.60<sup>a</sup> | 16.1<sup>a</sup> | 5.03<sup>b</sup> | 5.03<sup>abc</sup> | 0.44<sup>c</sup> |
| T8 | 93.2<sup>ab</sup> | 6.72□ | 20.72<sup>ab</sup> | 146.3<sup>a</sup> | 25.72<sup>ab</sup> | 12.59<sup>ab</sup> | 14.1<sup>b</sup> | 5.14<sup>b</sup> | 6.41<sup>a</sup> | 0.48<sup>abc</sup> |
| Irrigation |  |  |  |  |  |  |  |  |  |  |
| CF (I1) | 95.7 | 6.22 | 20.7 | 155.5 | 19.1 | 9.26 | 15.68 | 5.79 | 4.92 | 0.49 |
| DI (I2) | 92.0 | 5.85 | 20.3 | 156.8 | 24.3 | 12.47 | 15.20 | 5.19 | 5.75 | 0.46 |
| Fertilisers |  |  |  |  |  |  |  |  |  |  |
| TLL-RDF |  |  |  |  |  |  |  |  |  |  |
| (F1) | 93.2 | 23.37 | 20.1 | 152.9 | 17.9 | 9.51 | 15.62 | 5.63 | 5.39 | 0.48 |
| FP (F2) | 94.5 | 1.22 | 20.8 | 159.4 | 25.5 | 12.21 | 15.26 | 5.35 | 5.28 | 0.47 |
| Varieties |  |  |  |  |  |  |  |  |  |  |
| ADT-45<br>(V1) | 94.4 | 5.24 | 20.1 | 167.3 | 21.3 | 11.28 | 16.62 | 5.33 | 4.69 | 0.46 |
| BPT-5204<br>(V2) | 93.3 | 2.08 | 20.9 | 145.0 | 22.1 | 10.44 | 14.27 | 5.65 | 5.98 | 0.49 |
| Irrigation (I) | * | * | ns | ns | * | ** | * | ** | ** | * |
| Fertilisers<br>(F) | ns | ns | * | ns | ** | ** | * | * | ns | ns |
| Varieties (V) | ns | ** | * | ** | ns | ns | ** | * | ** | * |
| V x I | ns | ns | ns | ns | ns | ns | ns | ns | ns | ns |
| V x F x I | ns | ns | ns | ns | ns | ns | ns | ns | * | ns |
| F x I | * | ns | ns | ns | ns | ns | ns | ns | * | ns |

| Site 3 | Plant<br>height (in<br>cm) | Effective<br>tillers/plant | Panicle<br>length (in<br>cm) | Grain/panicle | Sterile<br>spikelet (in<br>numbers) | Unfilled<br>grains (%) | Test<br>weight<br>(1,000 | Grain yield<br>(in<br>tonnes/ha) | Straw yield<br>(in<br>tonnes/ha) | Harvest<br>index |
| --- | --- | --- | --- | --- | --- | --- | --- | --- | --- | --- |
|  |  |  |  |  |  |  | seeds) |  |  |  |
| T1 | 95.0<sup>ab</sup> | 6.78□□ | 19.75<sup>b</sup> | 171.3<sup>a</sup> | 11.16<sup>d</sup> | 6.24<sup>b</sup> | 17.28<sup>a</sup> | 5.64<sup>ab</sup> | 5.61<sup>b</sup> | 0.46<sup>ab</sup> |
| T2 | 97.8<sup>ab</sup> | 7.50□ | 20.42<sup>ab</sup> | 152<sup>a</sup> | 14.94<sup>cd</sup> | 7.87<sup>ab</sup> | 14.47<sup>bc</sup> | 6.10<sup>a</sup> | 7.73<sup>a</sup> | 0.53<sup>a</sup> |
| T3 | 94.5<sup>ab</sup> | 6.22□□ | 20.33<sup>ab</sup> | 170.4<sup>a</sup> | 19.42<sup>abc</sup> | 6.11<sup>b</sup> | 16.82<sup>a</sup> | 5.36<sup>ab</sup> | 5.48<sup>b</sup> | 0.44<sup>b</sup> |
| T4 | 103.3<sup>a</sup> | 6.67□□ | 20.87<sup>a</sup> | 158.7<sup>a</sup> | 23.06<sup>ab</sup> | 10.32<sup>ab</sup> | 13.95<sup>c</sup> | 5.66<sup>ab</sup> | 6.99<sup>ab</sup> | 0.47<sup>ab</sup> |
| T5 | 89.1<sup>b</sup> | 6.22□□ | 19.77<sup>b</sup> | 159.3<sup>a</sup> | 15.93<sup>bcd</sup> | 7.34<sup>ab</sup> | 16.82<sup>a</sup> | 5.17<sup>ab</sup> | 5.43<sup>b</sup> | 0.48<sup>ab</sup> |
| T6 | 100.2<sup>ab</sup> | 6.44□□ | 21.38<sup>a</sup> | 133<sup>a</sup> | 19.56<sup>abc</sup> | 10.32<sup>ab</sup> | 13.73<sup>c</sup> | 5.39<sup>ab</sup> | 6.01<sup>b</sup> | 0.46<sup>ab</sup> |
| T7 | 95.5<sup>ab</sup> | 5.67□ | 20.01<sup>ab</sup> | 159.6<sup>a</sup> | 21.32<sup>abc</sup> | 11.26<sup>a</sup> | 16.1<sup>ab</sup> | 4.94<sup>b</sup> | 5.28<sup>b</sup> | 0.43<sup>b</sup> |
| T8 | 99.7<sup>ab</sup> | 6.67□□ | 21.22<sup>a</sup> | 148<sup>a</sup> | 24.15<sup>a</sup> | 11.06<sup>a</sup> | 13.85<sup>c</sup> | 5.17<sup>ab</sup> | 6.29<sup>ab</sup> | 0.44<sup>b</sup> |
| Irrigation |  |  |  |  |  |  |  |  |  |  |
| CF (I1) | 97.6 | 6.74 | 20.34 | 163.1 | 17.1 | 7.64 | 15.6 | 5.69 | 6.46 | 0.48 |
| DI (I2) | 96.1 | 6.25 | 20.61 | 150.1 | 20.2 | 10.00 | 15.1 | 5.17 | 5.75 | 0.46 |
| Fertilisers |  |  |  |  |  |  |  |  |  |  |
| TLL-RDF |  |  |  |  |  |  |  |  |  |  |
| (F1) | 95.5 | 6.02 | 20.33 | 153.9 | 15.4 | 7.95 | 15.6 | 5.58 | 6.19 | 0.49 |
| FP (F2) | 98.3 | 3.13 | 20.62 | 159.3 | 22.0 | 9.69 | 15.2 | 5.28 | 6.01 | 0.45 |
| Varieties |  |  |  |  |  |  |  |  |  |  |
| ADT-45 | 93.5 | 4.95 | 19.98 | 165.2 | 17.0 | 7.74 | 16.8 | 5.28 | 5.45 | 0.46 |
| (V1) |  |  |  |  |  |  |  |  |  |  |
| BPT-5204 |  |  |  |  |  |  |  |  |  |  |
| (V2) | 100.2 | 0.26 | 20.97 | 148.0 | 20.4 | 9.89 | 14.0 | 5.58 | 6.76 | 0.48 |
| *Irrigation (I)* | *ns* | * | *ns* | *ns* | * | ** | * | ** | ** | *ns* |
| *Fertilisers (F)* | *ns* | *ns* | *ns* | *ns* | ** | * | *ns* | * | *ns* | ** |
| *Varieties (V)* | ** | * | ** | * | ** | ** | ** | * | ** | * |
| *V x I* | *ns* | *ns* | *ns* | *ns* | *ns* | *ns* | *ns* | *ns* | *ns* | * |
| *V x F x I* | *ns* | *ns* | *ns* | *ns* | *ns* | *ns* | *ns* | *ns* | *ns* | *ns* |
| *F x I* | *ns* | *ns* | *ns* | *ns* | *ns* | * | *ns* | *ns* | *ns* | *ns* |

| Site 4 | Plant height (in cm) | Effective tillers/plant | Panicle length (in cm) | Grain/panicle | Sterile spikelet (in numbers) | Unfilled grains (%) | Test weight (1,000 seeds) | Grain yield (in tonnes/ha) | Straw yield (in tonnes/ha) | Harvest index |
| --- | --- | --- | --- | --- | --- | --- | --- | --- | --- | --- |
| T1 | 89.3<sup>abc</sup> | 8.11 □ □ | 19.6<sup>ab</sup> | 172.7<sup>a</sup> | 13.49<sup>c</sup> | 8.01<sup>e</sup> | 16.97<sup>a</sup> | 5.6<sup>ab</sup> | 3.38<sup>bc</sup> | 0.50<sup>ab</sup> |
| T2 | 93.0<sup>ab</sup> | 9.44 □ | 20.8<sup>a</sup> | 154<sup>a</sup> | 17.97<sup>bc</sup> | 7.53<sup>de</sup> | 14.67<sup>bc</sup> | 6.0<sup>a</sup> | 5.37<sup>a</sup> | 0.53<sup>a</sup> |
| T3 | 100.2<sup>a</sup> | 9.06 □ | 19.6<sup>ab</sup> | 165.1<sup>a</sup> | 21.13<sup>b</sup> | 10.67<sup>cd</sup> | 17.22<sup>a</sup> | 5.53<sup>ab</sup> | 4.59<sup>ab</sup> | 0.48<sup>b</sup> |
| T4 | 102<sup>a</sup> | 9.67 □ | 21.3<sup>a</sup> | 144.6<sup>a</sup> | 23.06<sup>ab</sup> | 10.34<sup>cd</sup> | 14.68<sup>bc</sup> | 6.14<sup>a</sup> | 5.34<sup>a</sup> | 0.51<sup>ab</sup> |
| T5 | 88.5<sup>bc</sup> | 7.67 □ | 18.4<sup>b</sup> | 172.5<sup>a</sup> | 20.24<sup>b</sup> | 10.93<sup>bc</sup> | 16.22<sup>ab</sup> | 5.14<sup>b</sup> | 2.96<sup>c</sup> | 0.47<sup>b</sup> |
| T6 | 92.0<sup>abc</sup> | 8.44 □ □ | 19.6<sup>ab</sup> | 151.7<sup>a</sup> | 20.63<sup>b</sup> | 11.14<sup>bc</sup> | 14.06<sup>c</sup> | 5.78<sup>ab</sup> | 5.32<sup>a</sup> | 0.50<sup>ab</sup> |
| T7 | 83.8<sup>bc</sup> | 7.78 □ | 19.5<sup>ab</sup> | 154.3<sup>a</sup> | 26.85<sup>a</sup> | 14.51<sup>ab</sup> | 16.48<sup>ab</sup> | 5.28<sup>b</sup> | 3.02<sup>c</sup> | 0.45<sup>b</sup> |
| T8 | 83.0<sup>bc</sup> | 8.67 □ □ | 20.6<sup>ab</sup> | 142.1<sup>a</sup> | 27.88<sup>a</sup> | 16.41<sup>a</sup> | 14.87<sup>bc</sup> | 5.72<sup>ab</sup> | 5.43<sup>a</sup> | 0.51<sup>ab</sup> |
| Irrigation |  |  |  |  |  |  |  |  |  |  |
| CF (I1) | 96.1 | 9.07 | 20.3 | 159.1 | 19.0 | 9.14 | 15.9 | 5.82 | 4.67 | 0.51 |
| DI (I2) | 86.8 | 8.14 | 19.5 | 155.2 | 24.1 | 13.25 | 15.4 | 5.48 | 4.18 | 0.49 |
| Fertilisers |  |  |  |  |  |  |  |  |  |  |
| TLL-RDF |  |  |  |  |  |  |  |  |  |  |
| (F1) | 90.7 | 4.64 | 19.6 | 162.8 | 18.3 | 9.40 | 15.5 | 5.63 | 4.26 | 0.50 |
| FP (F2) | 92.3 | 0.80 | 20.3 | 151.5 | 24.8 | 12.98 | 15.8 | 5.67 | 4.60 | 0.49 |
| Varieties |  |  |  |  |  |  |  |  |  |  |
| ADT-45 |  |  |  |  |  |  |  |  |  |  |
| (V1) | 90.4 | 4.93 | 19.3 | 166.2 | 20.6 | 11.03 | 16.7 | 5.39 | 3.49 | 0.48 |
| BPT-5204 | 92.5 | 0.13 | 20.6 | 148.1 | 22.5 | 11.36 | 14.6 | 5.91 | 5.37 | 0.51 |
| (V2) |  |  |  |  |  |  |  |  |  |  |
| *Irrigation (I)* | ** | * | ** | *ns* | * | ** | ** | * | * | * |
| *Fertilisers (F)* | *ns* | *ns* | *ns* | *ns* | ** | ** | *ns* | *ns* | *ns* | *ns* |
| *Varieties (V)* | *ns* | * | ** | * | *ns* | *ns* | ** | ** | ** | ** |
| *V x I* | *ns* | *ns* | *ns* | *ns* | *ns* | *ns* | *ns* | *ns* | ** | *ns* |
| *V x F x I* |  |  |  |  |  |  |  |  |  |  |
| *F x I* |  |  |  |  |  |  |  |  |  |  |

| Site 5 | Plant height (in cm) | Effective tillers/plant | Panicle length (in cm) | Grain/panicle | Sterile spikelet (in numbers) | Unfilled grains (%) | Test weight (1,000 seeds) | Grain yield (in tonnes/ha) | Straw yield (in tonnes/ha) | Harvest index |
| --- | --- | --- | --- | --- | --- | --- | --- | --- | --- | --- |
| T1 | 80.0<sup>bc</sup> | 7.00□□ | 18.8<sup>bc</sup> | 161.4<sup>a</sup> | 12.4<sup>c</sup> | 15.9<sup>c</sup> | 16.1<sup>a</sup> | 4.59<sup>ab</sup> | 4.7<sup>ab</sup> | 0.48<sup>ab</sup> |
| T2 | 78.2<sup>bcd</sup> | 7.56□□ | 18.3<sup>c</sup> | 139.1<sup>a</sup> | 19.7<sup>bc</sup> | 16.1<sup>bc</sup> | 13.3<sup>c</sup> | 5.16<sup>a</sup> | 5.2<sup>a</sup> | 0.51<sup>a</sup> |
| T3 | 76.5<sup>cd</sup> | 8.22□□ | 18.1<sup>c</sup> | 156.3<sup>a</sup> | 15.6<sup>bc</sup> | 16.7<sup>bc</sup> | 16.08<sup>a</sup> | 4.01<sup>bc</sup> | 4.1<sup>abc</sup> | 0.47<sup>ab</sup> |
| T4 | 86.3<sup>ab</sup> | 9.56□ | 19.7<sup>abc</sup> | 147.7<sup>a</sup> | 24.6<sup>ab</sup> | 20.1<sup>ab</sup> | 13.4<sup>bc</sup> | 4.21<sup>bc</sup> | 4.8<sup>ab</sup> | 0.49<sup>ab</sup> |
| T5 | 84.3<sup>abc</sup> | 6.78□□ | 18.7<sup>bc</sup> | 162.1<sup>a</sup> | 20.1<sup>ab</sup> | 20.6<sup>ab</sup> | 14.9<sup>ab</sup> | 3.99<sup>c</sup> | 3.8<sup>c</sup> | 0.45<sup>b</sup> |
| T6 | 91.33<sup>a</sup> | 7.33□□ | 21.4<sup>a</sup> | 131.6<sup>a</sup> | 22.1<sup>ab</sup> | 24.6<sup>a</sup> | 13.4<sup>bc</sup> | 4.53<sup>ab</sup> | 4.5<sup>ab</sup> | 0.49<sup>ab</sup> |
| T7 | 85.33<sup>abc</sup> | 6.56□ | 19.6<sup>abc</sup> | 147.5<sup>a</sup> | 22.8<sup>ab</sup> | 17.7<sup>bc</sup> | 15.1<sup>ab</sup> | 4.03<sup>bc</sup> | 3.9<sup>bc</sup> | 0.44<sup>b</sup> |
| T8 | 86.2<sup>ab</sup> | 8.44□□ | 20.3<sup>ab</sup> | 133.7<sup>a</sup> | 30.6<sup>a</sup> | 20.9<sup>ab</sup> | 13.5<sup>bc</sup> | 4.21<sup>bc</sup> | 4.3<sup>abc</sup> | 0.49<sup>ab</sup> |
| Irrigation |  |  |  |  |  |  |  |  |  |  |
| CF (I1) | 80.25 | 8.08 | 18.72 | 151.1 | 18.1 | 17.2 | 14.7 | 4.49 | 4.68 | 0.49 |
| DI (I2) | 86.77 | 7.28 | 20.03 | 143.8 | 23.9 | 21.0 | 14.2 | 4.16 | 4.13 | 0.47 |
| Fertilisers |  |  |  |  |  |  |  |  |  |  |
| TLL-RDF |  |  |  |  |  |  |  |  |  |  |
| (F1) | 83.44 | 11.02 | 19.32 | 148.6 | 18.6 | 19.3 | 14.4 | 4.57 | 4.54 | 0.48 |
| FP (F2) | 83.58 | 9.92 | 19.43 | 146.3 | 23.4 | 18.9 | 14.5 | 4.09 | 4.27 | 0.47 |
| Varieties |  |  |  |  |  |  |  |  |  |  |
| ADT-45 |  |  |  |  |  |  |  |  |  |  |
| (V1) | 81.52 | 6.09 | 18.82 | 156.9 | 17.8 | 17.7 | 15.5 | 4.16 | 4.12 | 0.46 |
| BPT-5204 |  |  |  |  |  |  |  |  |  |  |
| (V2) | 85.50 | 2.62 | 19.93 | 138.0 | 24.3 | 20.4 | 13.4 | 4.50 | 4.69 | 0.49 |
| *Irrigation (I)* | * | * | ** | *ns* | ** | * | * | * | * | * |
| *Fertilisers (F)* | *ns* | *\*\** | *ns* | *ns* | *\** | *ns* | *ns* | *\*\** | *ns* | *ns* |
| *Varieties (V)* | *ns* | *\*\** | *\*\** | *\** | *\*\** | *ns* | *\*\** | *\** | *\** | *\*\** |
| *V x I* | *ns* | *ns* | *ns* | *ns* | *ns* | *ns* | *\** | *ns* | *ns* | *ns* |
| *V x F x I* |  |  |  |  |  |  |  |  |  |  |
| *F x I* |  |  |  |  |  |  |  |  |  |  |

*\*- Significant at 5%, \*\*- Significant at 1%, ns-nonsignificant. In a column, figures having the same letter(s) do not differ significantly, whereas figures with different letter(s) differ significantly, as per LSD at 5% level of significance. (T1-CF+V1+F1, T2-CF+V2+F1, T3-CF+V1+F2, T4-CF+V2+F2, T5-DI+V1+F1, T6-DI+V2+F1, T7 DI+V1+F2, T8 – DI+V2+F2)*

**Supplemental Table 3:** Water Usage under continuous and Drip irrigation (m^3^ha^-1^)

| Sites | Flood ($\text{m}^3\text{ha}^{-1}$) | Drip ($\text{m}^3\text{ha}^{-1}$) | Absolute Saving<br>($\text{m}^3\text{ha}^{-1}$) | Reduction (%) |
| --- | --- | --- | --- | --- |
| Site-1 | 12523 | 7668 | 4,855 | 38.80% |
| Site-2 | 14462 | 7887 | 6,575 | 45.50% |
| Site-3 | 15312 | 9357 | 5,955 | 38.90% |
| Site-4 | 13178 | 7678 | 5,500 | 41.70% |
| Site-5 | 16503 | 9508 | 6,995 | 42.40% |

**Supplemental Table 4:**
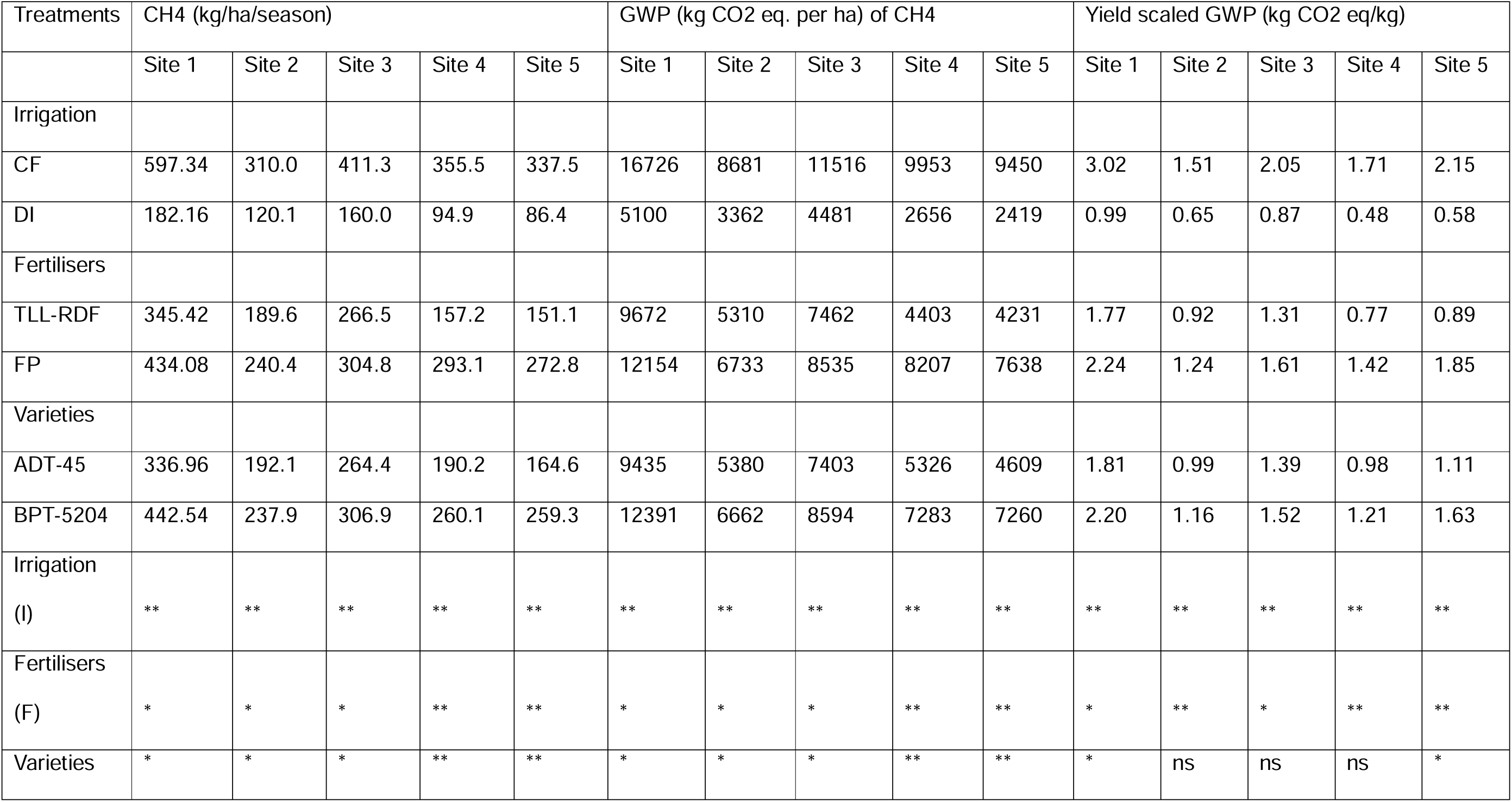
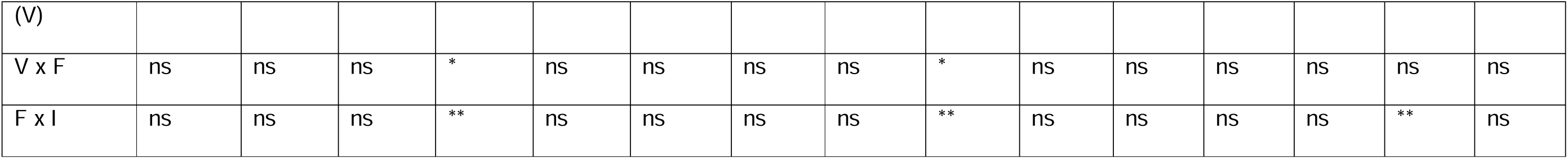
Methane emission, GWP and Yields scaled GWP of two rice varieties under two irrigations and two fertilizer combinations.

**Supplemental Figure 1:**
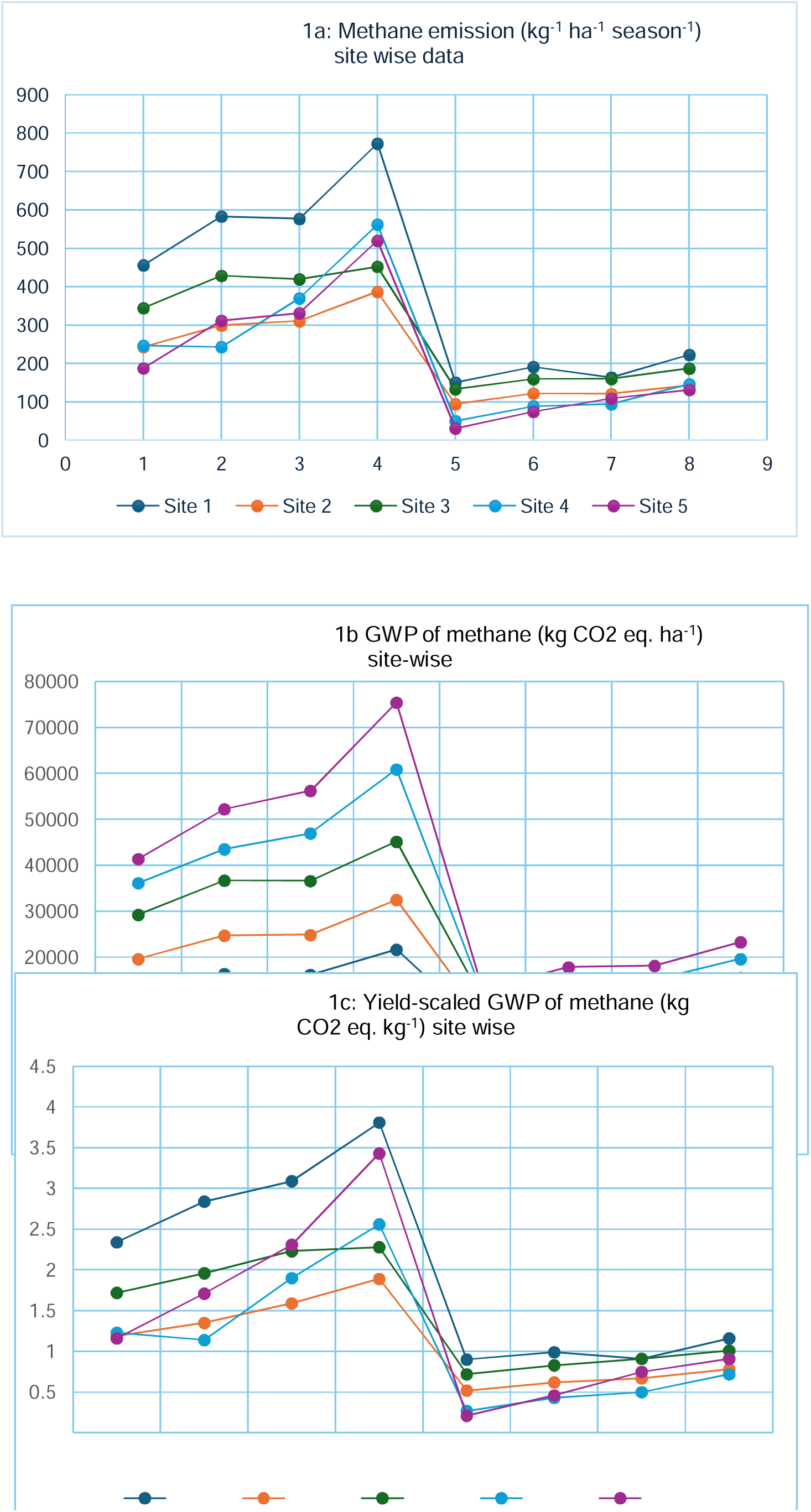
Site-wise data for methane emission, global warming potential (GWP) and yield-scaled global warming potential for eight treatments carried out in five sites in Sathyamangalam, Tamil Nadu, India

